# Motor Sequence Learning in Children and Adults: Age Differences in the Time Course of Brain Activation and Representational Stability

**DOI:** 10.64898/2026.05.12.724531

**Authors:** Maike Hille, Elisabeth Wenger, Eleftheria Papadaki, Yana Fandakova

**Affiliations:** Center for Lifespan Psychology, Max Planck Institute for Human Development, Lentzeallee 94, 14195 Berlin, Germany; Center for Environmental Neuroscience, Max Planck Institute for Human Development, Lentzeallee 94, 14195 Berlin, Germany; Institute for Mind, Brain and Behavior, HMU Health and Medical University, Olympischer Weg 1, 14471 Potsdam, Germany; Department of Psychology, University of Trier, Universitaetsring 15, 54296, Trier, Germany

## Abstract

Humans possess an astounding ability to acquire complex movement sequences with limited practice. Motor sequence learning engages a distributed network of brain regions that show distinct learning-related changes, often characterized by predominant involvement of the prefrontal cortex (PFC) early in learning and increasing engagement of the primary motor cortex (M1) later in learning. Because motor regions mature relatively earlier than the PFC during development, we examined how children and adults differ in the time course of neural changes underlying motor sequence learning. Using functional magnetic resonance imaging (fMRI), we compared brain activation in children (7–10 years, N = 39, 17 female) and adults (20–32 years, N = 39, 19 female) during an associative visuomotor learning task. In both age groups, response times decreased with sequence repetition, with greater reductions in adults than in children. Across age groups, early learning was associated with heightened PFC activation, whereas later learning was characterized by increased activation in left M1 and bilateral supplementary motor area. Children and adults showed comparable decreases in PFC activation and PFC–M1 connectivity with sequence repetition. In contrast, adults exhibited larger learning-related increases in activation and stability of multivariate patterns in left M1. Together, these findings indicate that although both age groups engaged the PFC similarly to support increased control demands in early learning, children showed less pronounced modulation of M1 activation and representational similarity, suggesting that M1’s capacity to form stable, sequence-related representations may still be developing in middle childhood.

## Introduction

In daily life, we perform a variety of motor actions such as typing, playing the piano, or video gaming. These skills are acquired through repeated execution of relevant sequential movements, enabling us to eventually perform them effortlessly (Clegg et al., 1998). The cognitive and neural processes underlying motor skill acquisition are often investigated by instructing participants to execute motor responses (e.g., button presses) in a specific sequence following a cue or by having them determine the sequence through trial and error (Karuza et al., 2014). Typically, repeatedly executing the same sequence is associated with decreases in errors and in response times (Foerde & Poldrack, 2009).

In adults, the early phase of learning a new motor sequence is characterized by heightened activation in the prefrontal cortex (PFC), reflecting increased cognitive control demands as new strategies and routines are established (Chein & Schneider, 2012; Halsband & Lange, 2006; Jenkins et al., 1994; Sakai et al., 1998). As sequences become well-learned, activation in cognitive control regions decreases (Foerde & Poldrack, 2009; Toni et al., 1998). The late phases of motor sequence learning are typically dominated by activation in primary motor cortex (M1) and supplementary motor area (SMA) (Dayan & Cohen, 2011; Toni et al., 1998) reflecting increased motor precision and more automatic task execution (Chein & Schneider, 2012; Halsband & Lange, 2006).

Learning-related changes are not limited to regional activation, but also involve changes in interactions between brain regions and changes in neural representations (Bassett et al., 2015; Dhawale et al., 2017; Lindenberger & Lövdén, 2019; Wiestler & Diedrichsen, 2013). Using a discrete sequence-production task, Bassett et al. (2015) demonstrated that connectivity between frontal cognitive control hubs (such as the PFC) and motor regions decreased with practice, indicating that sensorimotor regions became more autonomous with learning. Focusing on multivariate patterns, Wiestler and Diedrichsen (2013) showed that motor skill learning leads to more distinct activation patterns in motor regions for different trained sequences. Within a model of brain plasticity, Lindenberger and Lövden (2019) proposed that the self-similarity or stability of neural activation patterns associated with a specific behavior should increase with practice. In the context of motor sequence learning, this would correspond to higher representational similarity of multivoxel patterns across repetitions of the same sequence.

The regions involved in motor sequence learning differ in their developmental trajectories: while sensorimotor regions mature relatively earlier, PFC continues to develop well into young adulthood (Casey et al., 2005; Gogtay et al., 2004; Sydnor et al., 2021). Ongoing PFC development may contribute to slower learning in children, especially during early phases when cognitive control demands are relatively high (Barnea-Goraly et al., 2005; Diamond, 2002). Although sensorimotor regions mature relatively earlier, it remains unclear whether children engage motor regions as effectively as adults to facilitate the fine-tuning and automatizations of movement patterns, especially during later learning phases (Dayan & Cohen, 2011; Karni et al., 1995). Initial evidence from a serial reaction time task (Thomas et al., 2004) showed superior learning in adults compared to children (7–11 years) accompanied by greater premotor cortex activation. In contrast, children exhibited greater putamen activation than adults. These age differences were present throughout the task and did not change from early to late learning. Thus, it remains unclear to what extent the trajectory of motor and PFC engagement differs by age. In the present study, we sought to examine age differences in learning-related changes in the engagement of frontal and motor regions, and their interactions, during the time course of motor sequence learning.

In an associative visuomotor learning task performed in the MRI scanner, children between 7 and 10 years and adults were presented with four squares corresponding to four response buttons; through trial and error, the participants learned sequences of varying length by repeatedly practicing them until they could perform each sequence without errors at least 17 times (cf. Toni et al., 1998). We examined how performance, univariate and multivariate brain activation, and brain connectivity changed with sequence repetition and differed between age groups. Both age groups were anticipated to increase in performance with sequence repetition. Replicating previous results with a similar paradigm (Toni et al., 1998), adults were expected to show activation decreases in PFC from early to late learning and activation increases in motor regions with repetition. We hypothesized that due to protracted PFC development, children would show lower PFC activation early in learning compared to adults, resulting in less pronounced PFC activation decreases with sequence repetition. Furthermore, we expected that earlier-maturing motor regions in children might show a similar learning-related increase in activation as in adults. Alternatively, children might rely more strongly on other (e.g., subcortical; Thomas et al., 2004) brain regions, especially early in learning. Finally, we conducted exploratory analyses to examine whether PFC–M1 functional connectivity and the similarity of activation patterns in M1 changed differently in children and adults with learning.

## Results

### Successful Motor Sequence Learning in Children and Adults

The present experiment was designed such that participants had to discover and learn a given motor sequence through trial and error. After two consecutive correct executions of the sequence, participants were required to repeat the sequence until a total of 15 correct sequence repetitions were executed (i.e., including any intermediate erroneous repetitions). Thus, we first checked that our experimental manipulation was successful by comparing the average error rate in the early phase (i.e., repetitions until and including the first two subsequent correct sequence executions) and the late phase (i.e., all subsequent repetitions) in both age groups, collapsing across short and long sequences (see Figure 1). Paired t-tests revealed that the error rate decreased significantly from the early to the late learning phase in children (*t*(38) = 22.0, *p* < 0.001, *d* = 3.52) and in adults (*t*(38) = 51.3, *p* < 0.001, *d* = 8.21).

**Figure 1.**
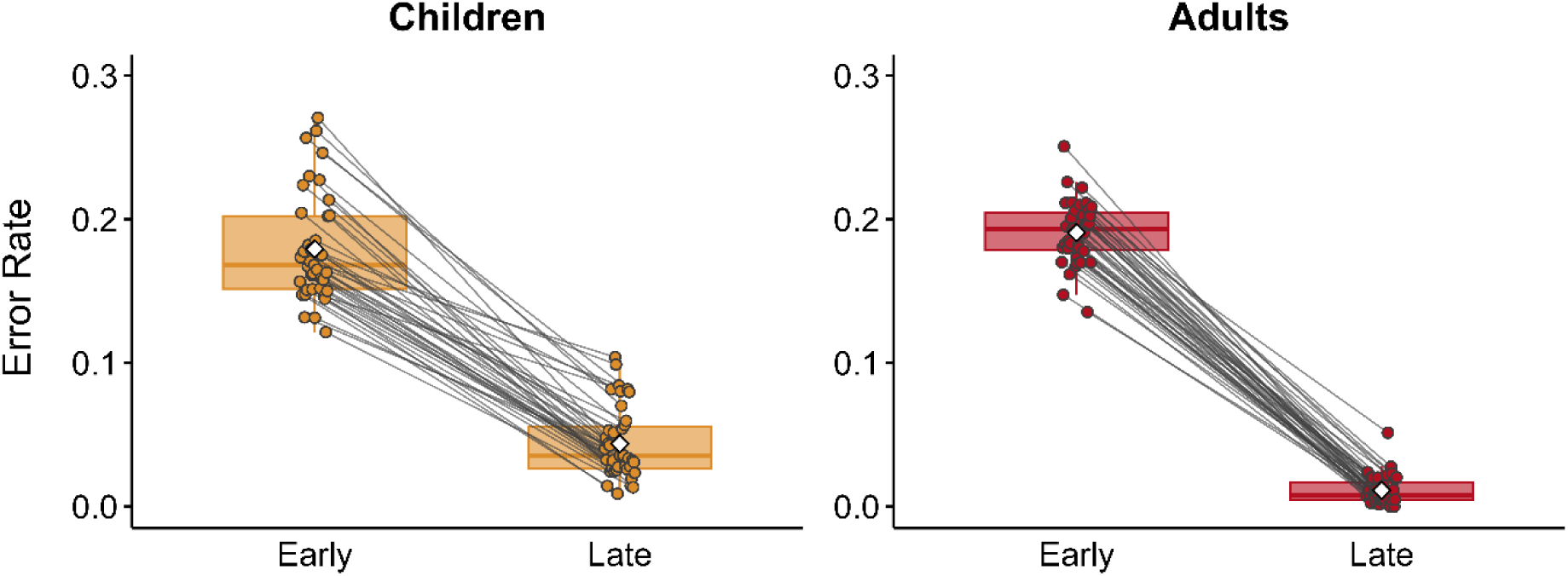
Error rate in early and late learning phases. Children are depicted in yellow and adults in red. Both age groups showed a significant decrease in the error rate from the early to the late learning phase.

In summary, with sequence repetition both age groups successfully learned the motor sequences, as evidenced by a significant reduction in execution errors over time.

### Learning-Related Improvements in RTs Across Children and Adults

Per experimental design, each participant repeated sequences of different length until each sequence was executed 17 times correctly, resulting in a varying number of sequence repetitions. To examine age differences in performance improvements across sequence repetitions, we first compared RTs between children and adults for the six-element sequences that were identical in both age groups.

#### Six-Element Sequences

RTs for the six-element sequences are illustrated in Figure 2. A linear mixed-effects model with log-transformed RTs as the dependent variable and age group (children vs. adults) and the linear and quadratic effects of sequence repetition as predictors revealed a significant negative linear effect of sequence repetition (*b* = -0.05, *SE* = 0.004, *p* < 0.001) along with a significant positive quadratic effect of sequence repetition (*b* = 0.002, *SE* = 0.0002, *p* < 0.001). These results indicate that across age groups RTs decreased quickly initially and slowed down with increasing sequence repetitions. There was also a significant effect of age group (*b* = 0.32, *SE* = 0.072, *p* < 0.001). Overall, children’s responses (*M* = 372 ms, *SD* = 144) were slower than adults’ (*M* = 232 ms, *SD* = 93.6). Additionally, we observed significant interactions between sequence repetition and age group (linear effect of sequence repetition × age group: *b* = 0.04, *SE* = 0.005, *p* < 0.001; quadratic effect of sequence repetition × age group: *b* = -0.001, *SE* = 0.0002, *p* < 0.001). These results indicate that while RTs decreased with sequence repetition in both age groups, adults showed greater initial linear RTs reductions than children, followed by more pronounced decelerations in later sequence repetitions. The very last, relatively sparsely sampled repetitions, showed a tendency for increasing RTs. Overall, results based on the six-element sequences suggest slower learning with sequence repetition in children compared to adults.

**Figure 2.**
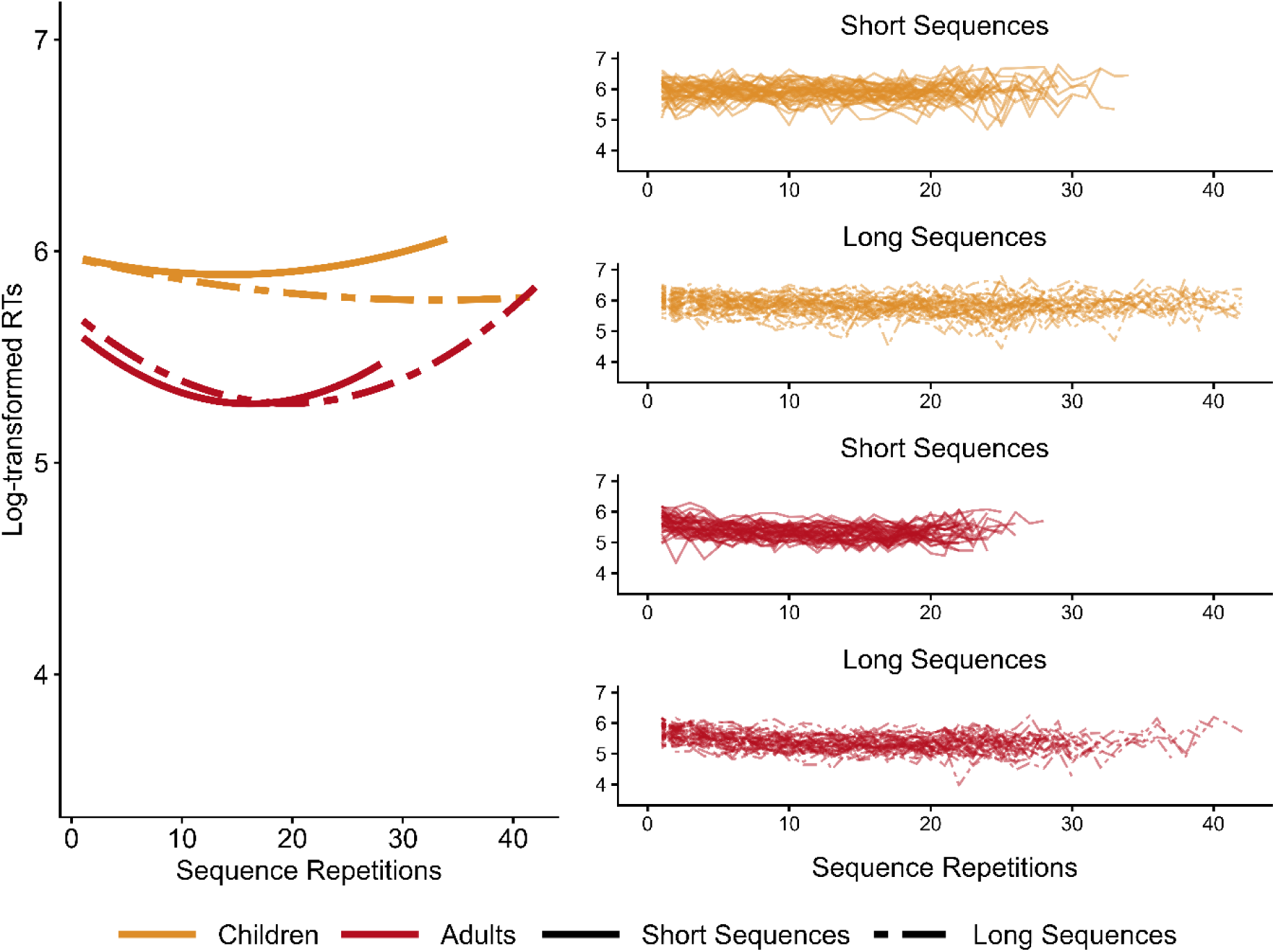
Log-transformed response times (RTs) across sequence repetitions. Children are shown in yellow and adults in red, with short sequences in solid lines and long sequences in dashed lines. The left panel depicts fixed-effect estimates from the linear mixed effect model testing for age differences in the linear and quadratic effects of sequence repetition. The right panel shows individual RT data. Both age groups showed faster RTs with sequence repetition, but the reduction was more pronounced in adults than in children.

#### Different Sequence Lengths

We next sought to examine how performance improvements differed between age groups for sequences of varying length that were matched in difficulty across age groups. For this purpose, we compared RT changes across sequence repetitions between age groups for short and long sequences. Short sequences contained 4 elements for children and 6 elements for adults, whereas long sequences contained 6 elements for children and 8 elements for adults. Thus, while the above analysis was based on a comparison of the exact same sequences between children and adults, we next compared sequences that were matched in difficulty between age groups.

We fitted a linear mixed-effects model with log-transformed RTs as the dependent variable. The model included the fixed effects of sequence repetition (modeled with linear and quadratic terms), age group (children vs. adults), and sequence length (short vs. long). The best fitting model showed significant main effects of sequence repetition (linear *b* = -0.04, *SE* = 0.002, *p* < 0.001; quadratic *b* = 0.001, *SE* = 0.0001, *p* < 0.001) and age group (*b* = 0.36, *SE* = 0.05, *p* < 0.001). Additionally, we found significant interactions between sequence repetition and age group (linear effect × age group: *b* = 0.03, *SE* = 0.003, *p* < 0.001; quadratic effect × age group: *b* = -0.001, *SE* = 0.0001, *p* < 0.001). These results echo those reported above. While both age groups improved with sequence repetition, these improvements were more pronounced in adults than in children and were accompanied by more pronounced slowing down during the last sequence repetitions (Figure 2). We also observed an interaction between age group and sequence length (*b* = -0.09, *SE* = 0.01, *p* < 0.001): across sequence repetitions, adults were overall faster for short (*M* = 232, *SD* = 93.6 ms) relative to long (*M* = 240 ms, *SD* = 98.2 ms) sequences, whereas children were faster for long sequences (*M* = 372 ms, *SD* = 144 ms) than for short sequences (*M* = 401 ms, *SD* = 166 ms). In addition, there was a significant interaction between the quadratic effect of sequence repetition and sequence length (*b* = -0.0002, *SE* = 0.00004, *p* < 0.001), reflecting the fact that RT increases towards the end of training were more likely for short than for long sequences. Note that these last repetitions were more sparsely sampled and not present for all participants (see Figure 2). The main effect of sequence length was not reliable (*b* = 0.08, *SE* = 0.06, *p* = 0.21). Model comparisons indicated that the three-way interactions including sequence repetition, age group, and sequence length were unwarranted (all *ps* > 0.05). Results did not change when we used raw RTs or inverse efficiency scores, which combine RTs with accuracy (Table S1–S4).

Taken together, across both short and long sequences, children and adults successfully learned the sequences with repetition as indicated by significant RT decreases. Adults showed more substantial improvements with sequence repetition than children. Next, we sought to test the degree to which these differences in motor sequence learning between children and adults were driven by differential engagement of the brain’s control and motor systems.

### Neural Underpinnings of Motor Sequence Learning Across Children and Adults

First, we investigated the brain regions supporting motor sequence learning during the early and late phases of learning. To this end, we performed a whole-brain analysis comparing the early versus late learning phases, collapsing across both age groups. Since our behavioral results revealed no distinguishable age group differences in performance improvements between short and long sequences, we collapsed across sequence lengths for these analyses. The contrast of early > late learning phase (*ppeak* < 0.05, FWE-corrected, *k* = 50 voxels) revealed enhanced activation across a broad frontoparietal network (Figure 3), including the lateral anterior PFC, dorsolateral PFC (dlPFC), inferior parietal lobule, superior parietal lobule, dorsal anterior cingulate cortex, anterior insula, bilateral caudate, inferior temporal gyrus, and precuneus. The opposite contrast of late > early learning phase (*ppeak* < 0.05, FWE-corrected, *k* = 50 voxels) demonstrated enhanced activation in the left M1, left and right SMA, medial PFC, left superior and middle temporal gyrus, and posterior cingulate cortex.

**Figure 3.**
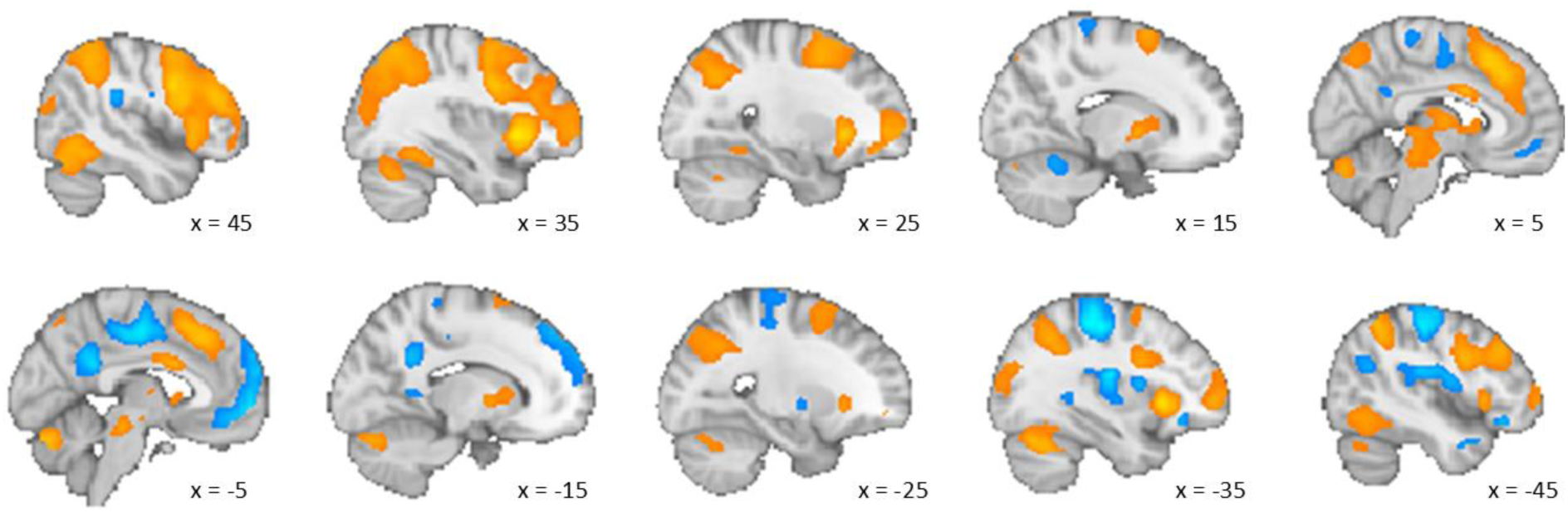
Brain activation associated with motor sequence learning. Brain activation of the early (in orange) and late (in blue) phases of learning motor sequences across both age groups and all sequence lengths (*ppeak* < 0.05, FWE-corrected, *k* = 50 voxels).

Whole-brain comparisons between children and adults (early vs. late learning phases) revealed no clusters surviving *ppeak* < 0.05, FWE-corrected, *k* = 50 voxels.

Taken together, across both children and adults, the early learning phase was characterized by enhanced activation in a broad network of lateral frontal and parietal regions, whereas the late learning phase entailed enhanced activation in motor, medial frontal, and temporal regions. Next, we sought to examine the degree to which these regions were differentially engaged across sequence repetitions between children and adults. Here we focused on brain regions previously implicated in motor skill learning during early (i.e., PFC) vs. late (i.e., M1, SMA) learning phases. For completeness, results for the left and right putamen ROIs, demonstrating overall enhanced activation in children than in adults (Figure S1 and Table S5, S6) are reported in detail in the Supplemental Material.

### Similar Decreases in PFC Activation Across Repetitions in Children and Adults

We first focused on the left and right IFG ROIs, which showed enhanced activation during early relative to late learning phases. To examine age-related differences in activation changes across repetitions, we predicted parameter estimates of each sequence repetition by the linear and quadratic effects of sequence repetition, age group (children vs. adults), and their interactions. Here, we focused on the first 17 sequence repetitions that were available for each participant in the present study.

In the left IFG (Figure 4A), replicating the whole-brain results above, we observed a significant negative linear decrease with sequence repetition (*b* = -0.49, *SE* = 0.05, *p* < 0.001) accompanied by a significant quadratic effect (*b* = 0.02, *SE* = 0.003, *p* < .001), indicating deceleration of activation decreases over time. The main effect of age group was not significant (*b* = 0.66, *SE* = 0.44, *p* = 0.14) and the interaction between sequence repetition and age group was not significant (*χ^2^* (2) = 4.01, *p* = 0.135). Results were similar in the right IFG (Figure 4B). Here, we also found a significant negative linear effect of sequence repetition (*b* = -0.49, *SE* = 0.04, *p* < 0.001) accompanied by a significant main quadratic effect (*b* = 0.02, *SE* = 0.002, *p* < 0.001), as expected based on the whole-brain results. The main effect of age group (*b* = -0.21, *SE* = 0.28, *p* = 0.46), and the interaction between sequence repetition and age group (*χ^2^* (2) = 2.24, *p* = 0.33) were not significant.

**Figure 4.**
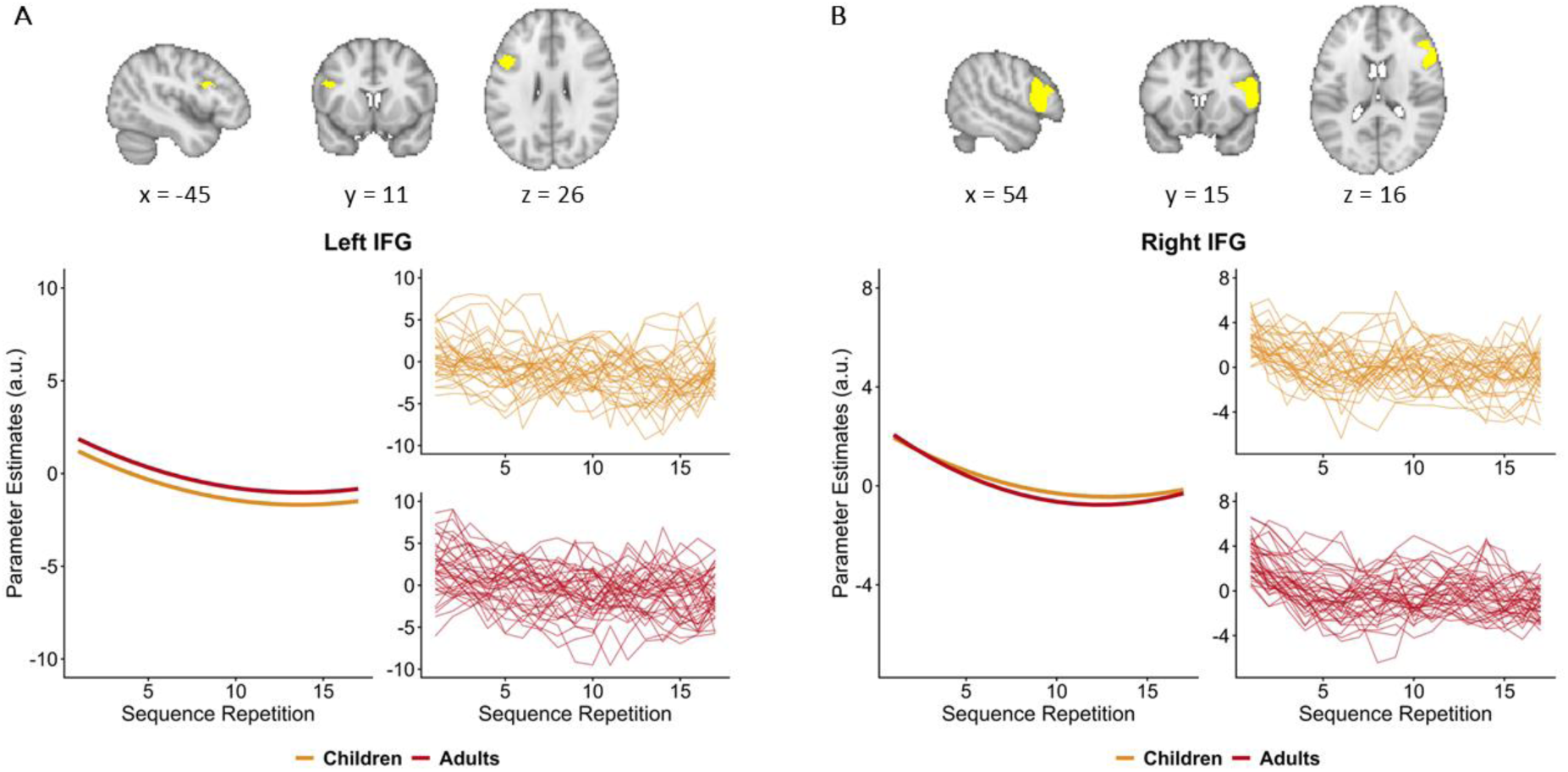
Brain activation in left and right IFG across sequence repetitions during learning. Model estimates for left (***A***) and right (***B***) IFG. For each ROI, the left panel depicts fixed-effect estimates from the linear mixed effect model testing for age-related differences in the linear and quadratic effects of sequence repetition. The right panel shows individual activation data for the first 17 sequence repetitions for children (in yellow) and adults (in red). Adults and children demonstrated similar decreases in activation with sequence repetition.

In summary, children and adults exhibited comparable decreases in PFC activation with repeated execution of the sequences.

### Age Differences in Left M1 Activation Increases Across Repetitions

We next focused on the left M1 and bilateral SMA ROIs, which exhibited increased activation during the late learning phase in the whole-brain analyses across age groups. Restricting our analysis to the first 17 sequence repetitions, we modeled the parameter estimates for each sequence repetition as a function of the linear and quadratic effects of sequence repetition, age group, and their interactions. In the left M1, we observed reliable linear (*b* = 0.43, *SE* = 0.05, *p* < 0.001) and quadratic (*b* = -0.02, *SE* = 0.003, *p* < 0.001) effects of sequence repetition. Recapitulating the whole-brain results above, in both age groups M1 activation initially increased, with increases slowing down over time (Figure 5A). Furthermore, we observed significant interactions between sequence repetition and age group (age group × linear effect of sequence repetition: *b* = 0.25, *SE* = 0.07, *p* = 0.001; age group × quadratic effect of sequence repetition: *b* = -0.01, *SE* = 0.004, *p* = 0.005), suggesting that the initial increase as well as the subsequent slowing down in M1 activation were more pronounced in adults than in children. The main effect of age group was not significant (*b* = -0.16, *SE* = 0.83, *p* = 0.85).

**Figure 5.**
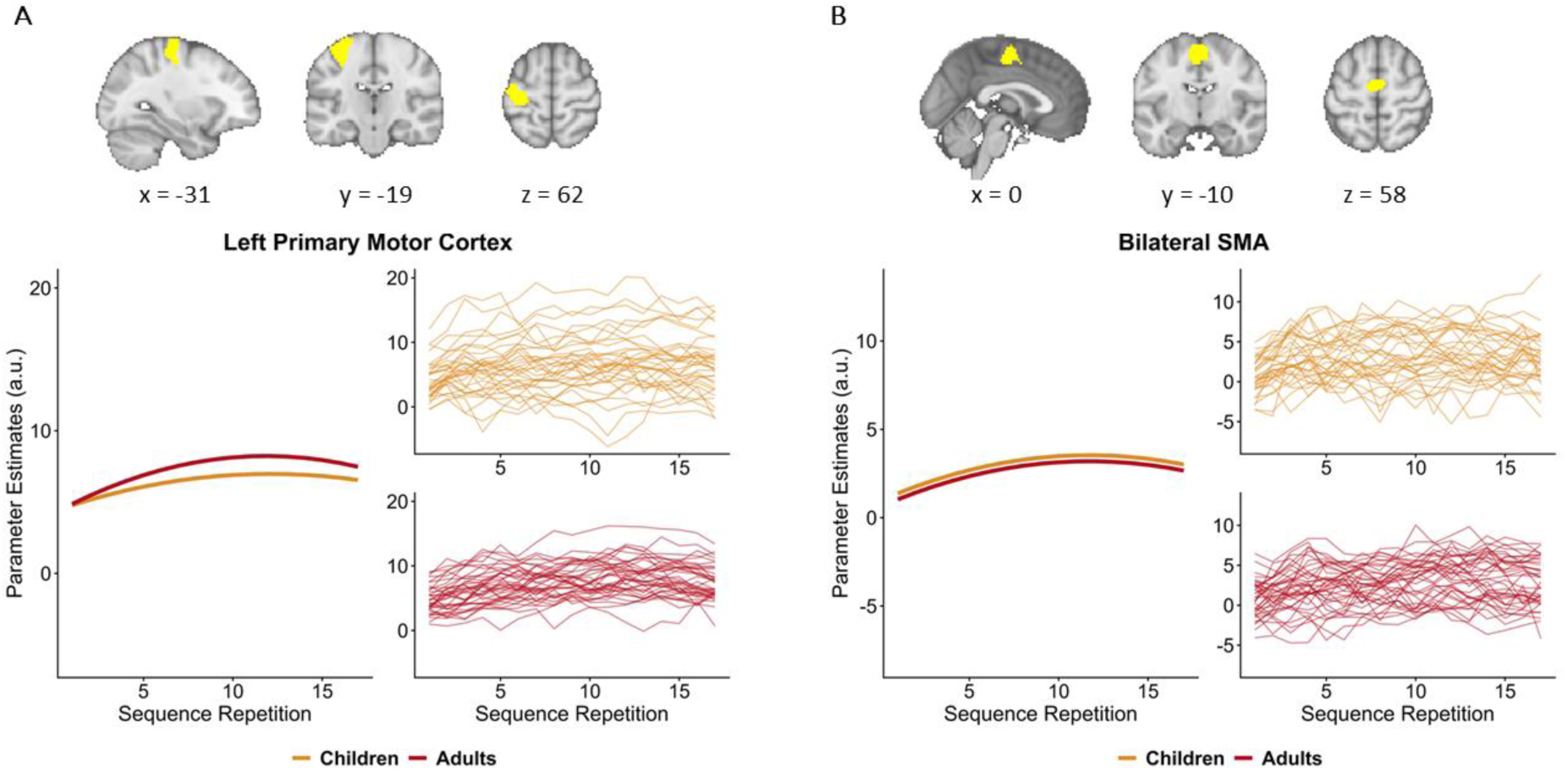
Brain activation in left M1 and bilaterial SMA across sequence repetitions during learning. Model estimates for left M1 (***A***) and bilateral SMA (***B***). For each ROI, the left panel depicts fixed-effect estimates from the linear mixed effect model testing for age differences in the linear and quadratic effects of sequence repetition. The respective right panel shows individual activation data for the first 17 sequence repetitions for children (in yellow) and adults (in red). In both regions, adults and children showed an increase in activation with sequence repetition followed by a decrease. In left M1, this pattern was more pronounced in adults than in children.

Similar results were observed in the bilateral SMA (Figure 5B). There was a significant positive linear effect of sequence repetition (*b* = 0.44, *SE* = 0.04, *p* < 0.001) as well as a significant quadratic effect (*b* = -0.02, *SE* = 0.002, *p* < 0.001), mirroring the whole-brain results reported above. The main effect of age group was not significant (*b* = -0.34, *SE* = 0.58, *p* = 0.56). Likelihood ratio tests showed that the interactions between age group and the linear effect of sequence repetition (*χ^2^*(1) = 3.47, *p* = 0.06) as well as between age group and the quadratic effect of sequence repetition (*χ^2^* (1) = 0.67, *p* = 0.41) did not significantly improve model fit.

In summary, across sequence repetition adults demonstrated more pronounced increases in activation in the left M1 than children.

Next, we examined how differences in brain activation were related to differences in task performance. For this purpose, we created two performance groups in each age group based on a median split of average RTs across all sequence repetitions (median = 232 s in adults; median = 404 s in children). We fitted a linear mixed-effects model predicting M1 parameter estimates, with performance group (high vs. low performance) as an additional factor to the ones above. The best-fitting model comprised the main effects of linear and quadratic sequence repetition, age group, and performance group along with interactions between the linear sequence repetition × age group × performance group and quadratic sequence repetition × age group. In this model, in line with the effects reported above, we found a significant interaction between linear sequence repetition × age group × performance group (*b* = -0.09, *SE* = 0.03, *p* = 0.005). All remaining effects involving performance group were not significant (all *ps* > 0.05). Follow-up models in each age group revealed that in adults, there was a significant interaction between linear sequence repetition × performance group (*b* = -0.1, *SE* = 0.02, *p* < 0.001) such that adults in the high-performance group exhibited a greater increase in M1 activation compared to those in the low-performance group (Figure 6). There were no differences in activation between high and low performers among children (*b* = -0.002, *SE* = 0.03, *p* = 0.94). Mixed-effects models revealed no significant differences in brain activation between performance groups in left IFG, right IFG, or bilateral SMA.

**Figure 6.**
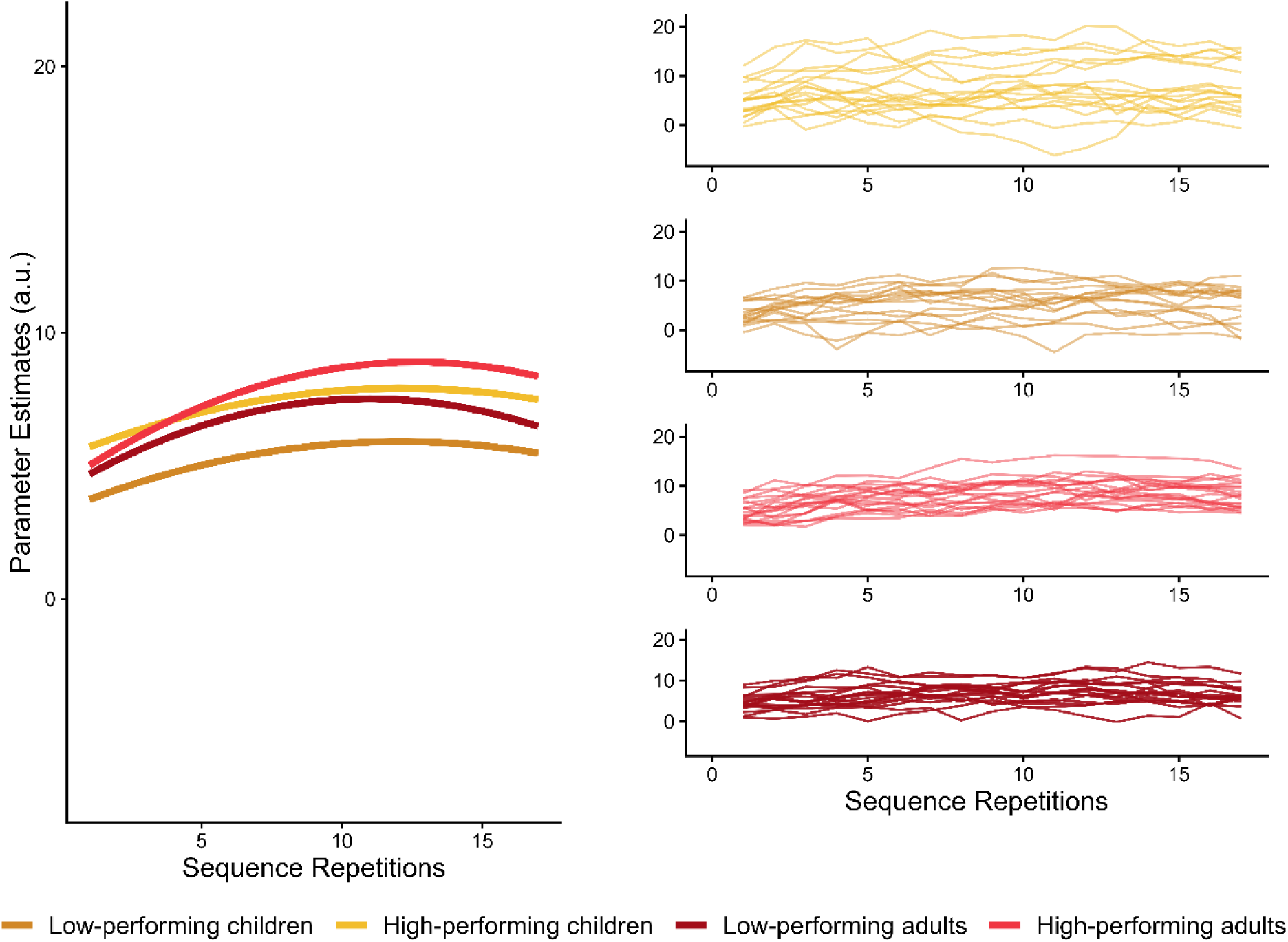
Performance-related differences in left M1 activation. Differences in brain activation in left M1 between performance groups (high vs. low performers) were evident in adults, but not in children. The left panel shows fixed-effect estimates from the linear mixed-effects model examining how brain activation varied across performance groups, considering its interactions with linear and quadratic sequence repetition and age group. The right panel shows individual M1 activation data for the first 17 sequence repetitions for children (in yellow) and adults (in red), split for low performers (darker color) and high performers (lighter color).

### Changes in Neural Activation Patterns in Left M1 Across Repetitions

As the preceding univariate analyses revealed differential effects of sequence repetition on left M1 activation between children and adults, we conducted an exploratory RSA to further characterize learning-related differences between the age groups. As learning progresses, we expected sequence representations to become more stable, resulting in increasing correlations between the activation patterns of two successive correct executions of the same sequence over time. Note that in contrast to the univariate analyses based on the first 17 sequence repetitions, we focused on the 17 correctly executed sequences that were available by design for each participant here. Z-transformed correlations between two subsequent correct sequence repetitions were predicted by the linear and quadratic effects of repetition pair, age group, and their interactions. The best fitting model indicated significant effects of repetition pair (linear: *b* = -0.015, *SE* = 0.005, *p* = 0.002; quadratic: *b* = 0.002, *SE* = 0.0003, *p* < 0.001). The estimated results are depicted in Figure 7 and demonstrate that pattern similarity initially decreased, and then increased with repetition pair. In addition, we found a significant interaction between age group and the linear effect of repetition pair (*b* = -0.008, *SE* = 0.0024, *p* = 0.001). Follow-up models in each age group indicated that while both children and adults displayed a reliable initial decrease in correlations (children: *b* = -0.024, *SE* = 0.008, *p* = 0.003; adults: *b* = -0.016, *SE* = 0.006, *p* = 0.021), the decrease was more pronounced in children. The interaction between age group and the quadratic effect of repetition pair was not significant (*χ^2^*(1) = 0.005, *p* = 0.946).

**Figure 7.**
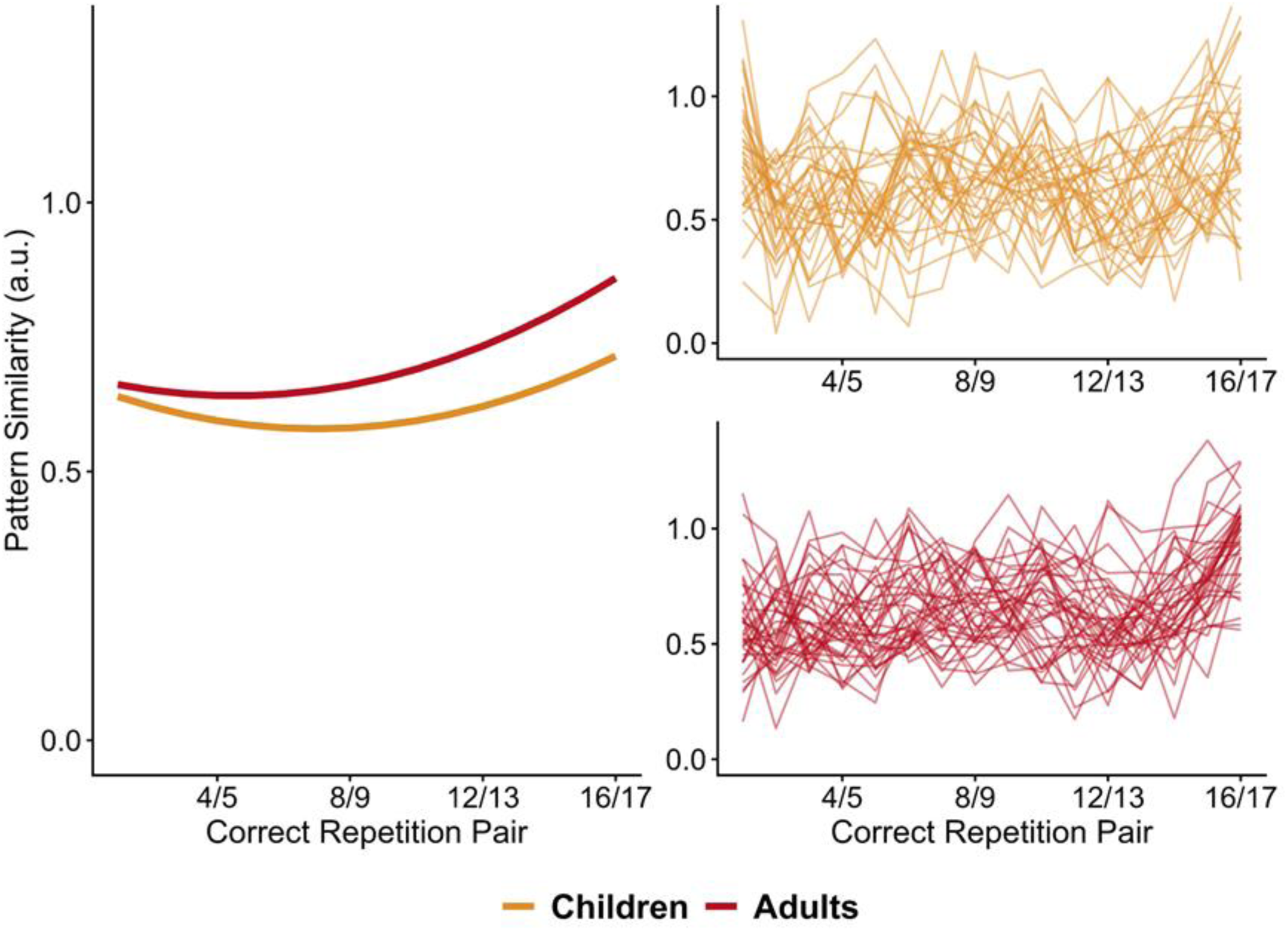
Similarity of neural activation patterns in the left M1. The left panel shows fixed-effect estimates from the linear mixed effect model testing for age differences in the linear and quadratic effects of repetition pair. Children’s data are shown in yellow and adults’ data in red. The right panel depicts individual pattern similarity data for the 16 sequence repetition pairs. While the initial decrease in representational similarity was greater in children than in adults, the subsequent increase was comparable between the two age groups.

Given this pattern of a greater initial decrease in children followed by a similar later increase in both groups, it is reasonable to expect group differences at the final repetition pair. Accordingly, a post-hoc comparison of age groups at the final repetition pair 16/17 revealed significantly higher pattern similarity in adults than in children (repetition pair 16/17: *b* = 0.15, *SE* = 0.03, *p* < 0.001). There were no significant differences in overall representational similarity between age groups (*b* = -0.015, *SE* = 0.031, *p* = 0.63).

Together, these results indicate that sequence learning in the left M1 was characterized by an initial decrease in representational similarity at the onset of learning, followed by increases in correlations later during learning. While the initial decrease in representation similarity was more pronounced in children than in adults, the increase in representation similarity in the late phases of learning was similar between children and adults, leading to greater representational similarity in adults than in children at the last sequence repetition pair.

### Changes in Functional Connectivity Between PFC and Left M1 Across Repetitions

Given the potential role of coordinated activation between the PFC and left M1 for motor sequence learning, we exploratorily examined how their functional connectivity changed with sequence repetition, focusing on the first 17 repetitions available for each participant. To do this, we predicted parameter estimates of pairwise connectivity by the linear and quadratic effects of sequence repetition and age group. We observed significant main effects of sequence repetition (linear: *b* = -0.018, *SE* = 0.006, *p* = 0.0027; quadratic: *b* = 0.0007, *SE* = 0.0003, *p* = 0.018), indicating that in both age groups, functional connectivity between PFC and left M1 decreased with sequence repetition, with less pronounced declines in later sequence repetitions (Figure 8). The main effect of age group was not significant (*b* = 0.005, *SE* = 0.026, *p* = 0.85) and the interactions between age group and sequence repetition (linear: *χ^2^*(1) = 2.47, *p* = 0.12; quadratic: *χ^2^*(1) = 1.24, *p* = 0.27) were unwarranted.

**Figure 8.**
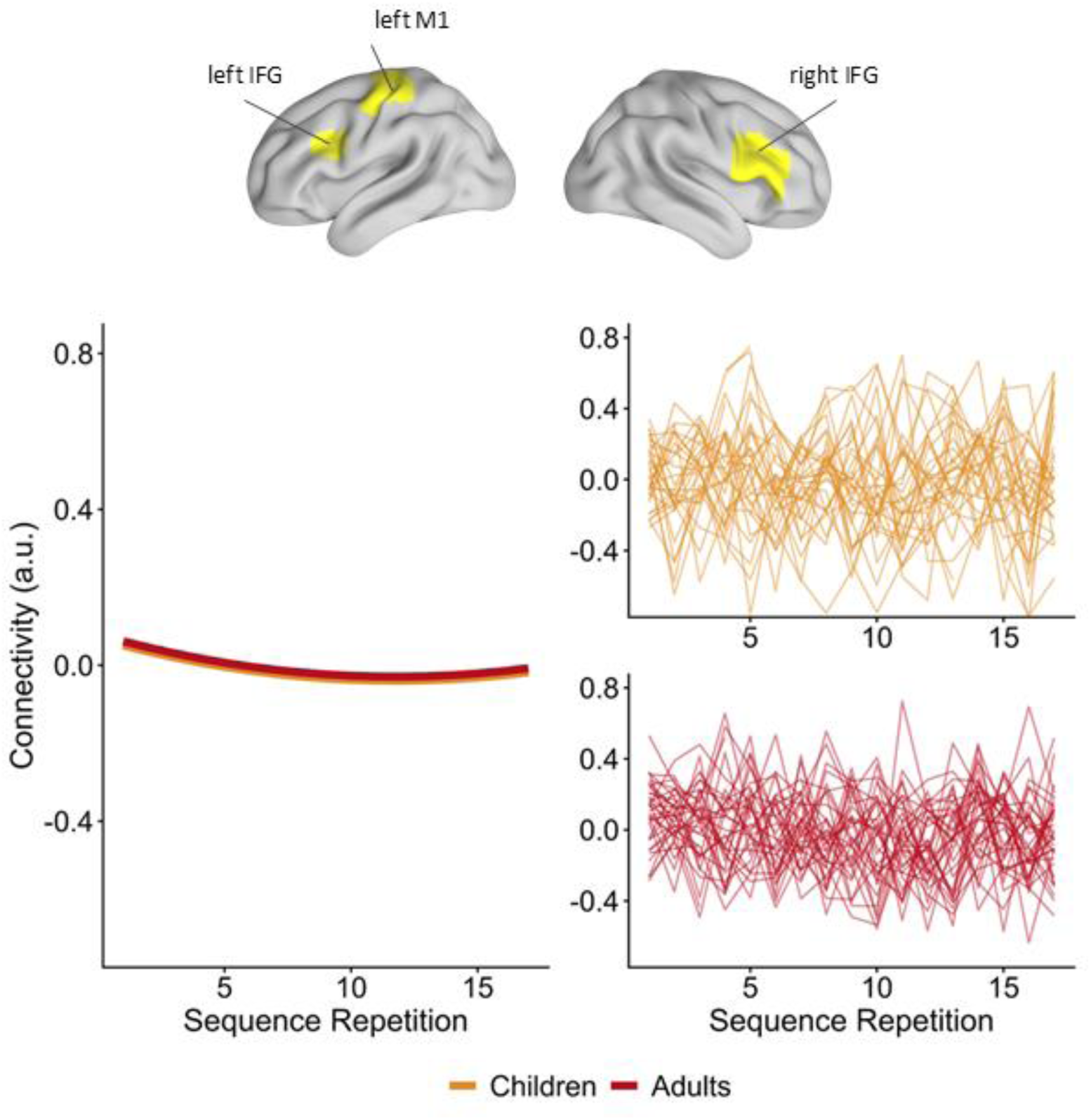
Functional connectivity between PFC (left & right IFG) and left M1 for the first 17 sequence repetitions. The left panel depicts fixed-effect estimates from the linear mixed effect model testing for age differences in the linear and quadratic effects of sequence repetition. The right panel shows individual connectivity data for the first 17 sequence repetitions for children (in yellow) and adults (in red). Functional connectivity between the PFC and left M1 decreased similarly in both age groups during learning.

In summary, the functional connectivity between the PFC and left M1 decreased during learning, and the decreases were comparable across age groups.

## Discussion

We investigated motor sequence learning across multiple motor sequences in 7–10-year-old children and adults to examine age differences in the neural processes supporting motor skill learning. Adults learned more efficiently than children, as indicated by steeper RT decreases with sequence repetition. Across age groups, the early learning phase was marked by heightened activation across a broad frontoparietal network, prominently including bilateral lateral PFC regions. In contrast, motor regions, including the left M1 and bilateral SMA, exhibited enhanced activation during the late learning phase. Across sequence repetitions, children and adults showed comparable decreases in PFC activation and in task-based PFC–M1 connectivity. Compared to adults, children showed less pronounced activation increases with sequence repetition in left M1. Furthermore, among adults, activation increases were more pronounced for high-performing than for low-performing individuals. Multivariate analyses corroborated the less efficient learning in children: both age groups showed increased similarity of activation patterns in left M1, but representational similarity was lower in children at the end of practice.

To date, relatively few studies have investigated age differences in motor sequence learning, typically focusing on variants of the serial reaction time task. Findings from these studies are mixed (Zwart et al., 2019): some reported no age differences in motor sequence learning, supporting a developmental invariance model (Karatekin et al., 2007; Mayor-Dubois et al., 2016; Meulemans et al., 1998), while others found age-related differences, with some studies observing more effective learning in children or adolescents than in adults (Fischer et al., 2007; Janacsek et al., 2012), and others reporting superior performance in adults compared to children (Jongbloed-Pereboom et al., 2019; Lukács & Kemény, 2015; Savion-Lemieux et al., 2009; Thomas et al., 2004). However, the exact trajectory remains unclear and appears to be mostly influenced by methodological differences, such as using raw or normalized RTs for analysis or the influence of intentional learning (Zwart et al., 2019).

Some evidence suggests that learning differs between incidental and intentional sequence learning with more pronounced age differences in performance and explicit sequence knowledge typically observed under explicit task conditions (Jongbloed-Pereboom et al., 2019; Karatekin et al., 2007; Nemeth et al., 2013; Weiermann & Meier, 2012). Importantly, in our paradigm, participants were explicitly asked to learn the motor sequences through trial and error. Thus, the observed RT changes are less likely to reflect differences in task predictability as typically assessed in serial reaction time tasks which compare performance of repeated sequences to random sequences. Our results are in line with Karatekin et al. (2007) who found that children (8–17 years) performed worse than adults under intentional learning conditions, whereas incidental learning yielded comparable performance across age groups. This pattern of results suggests that intentional sequence learning may place greater demands on strategic processes, possibly contributing to less effective motor learning in children (Hong et al., 2019; Karatekin et al., 2007). However, it should be noted that distinguishing between incidental and intentional learning processes is challenging, as both may contribute to procedural learning in overlapping and coordinated ways (Hong et al., 2019).

In all serial reaction time task studies above, the sequences were identical in length (usually 10 elements long) for all age groups. The present study extends those results by using difficulty-adapted sequences and showing that age differences in motor sequence learning persist. Children often display more variable performance than adults in perceptual learning as well, which has been attributed to less effective attention focusing (Moore et al., 2011) suggesting this as one possible contributing factor in the present study as well. Distributing attention during learning and information processing can lead to slower and less efficient performance on the task at hand due to the concurrent processing of irrelevant information (Chein & Schneider, 2012; Desimone et al., 1995). However, a broader attentional focus also enables children to learn more task-irrelevant information (Plebanek & Sloutsky, 2017; Tandoc et al., 2024) and may be an advantage in skill acquisition situations that require the exploration of different strategies (Fandakova & Wenger, 2024).

Our findings of less pronounced RT improvements in children align with the results reported by Thomas et al. (2004) who found more pronounced RT improvements in adults compared to 7–11-year-old children in a serial reaction time task. Similar trends were also observed by Savion-Lemieux et al. (2009) who investigated 6–10-year-old children and adults with a variant of the serial reaction time task.

At the neural level, we observed increased PFC activation during early learning, consistent with previous research on the progression of brain activation during motor learning in adults (Jenkins et al., 1994; Sakai et al., 1998; Toni et al., 1998; Wu et al., 2004). This finding aligns with the notion that control systems play a crucial role in the initial scaffolding of learning (Chein & Schneider, 2012; Hong et al., 2019; Koechlin et al., 2003). With practice, the observed decrease in PFC activation may reflect the formation of more specific task representations (Garner & Dux, 2015) or the use of more efficient strategies (Poldrack, 2000). As expected, regions recognized as representation systems, including M1 and SMA (Chein & Schneider, 2012; Dayan & Cohen, 2011), showed heightened activation during later phases, when the learning process is assumed to become increasingly more efficient (Doyon et al., 2018). The activation increase likely reflects these regions’ roles in integrating sensory and motor associations and storing relevant representations (Chein & Schneider, 2012; Penhune & Steele, 2012; Sanes & Donoghue, 2000). The temporal dissociation in PFC and M1 activation in the present study is closely aligned with a recent study that used inhibitory repetitive transcranial magnetic stimulation to investigate the causal roles of dlPFC and M1 during motor sequence learning (Nguyen et al., 2025). The results showed that transiently disrupting activity in these regions immediately prior to the task produced impairments in performance. Notably, dlPFC disruption more negatively affected performance on minimally trained sequences, whereas M1 disruption showed relatively larger behavioral deficits for highly trained sequences. This pattern appears to reflect a stronger reliance on frontal regions early in learning and a heavier reliance on motor regions later in learning. Note that, although we distinguish between relatively earlier and later phases of learning, the changes observed in our study would still be considered part of the initial phase of acquiring a new skill (Dayan & Cohen, 2011). As such, they are unlikely to reflect automatization, but rather the transformations that may be paving the way for later automatization. To better understand the link between early changes in brain activation and rapid structural plasticity (Olivo et al., 2022; Taubert et al., 2016) especially in light of developmental differences, future research should include repeated structural imaging within single learning sessions.

Of note, our results suggest that the temporally dissociable roles of PFC and M1 are already present in middle childhood and during very early learning, indicating that the dynamic engagement of control and representation systems represents a fundamental feature of motor sequence learning across development.

As learning progresses, domain-specific systems become more autonomous, showing reduced integration with control regions (Bassett et al., 2015) and decreased neural variability (Dhawale et al., 2017). These changes are thought to reflect the selection of the most efficient neural pathways (Bassett et al., 2015; Lindenberger & Lövdén, 2019). In line with this interpretation, both children and adults showed similar decreases in functional connectivity between PFC and left M1, as well as less variable, more stable neural representations in left M1 over time.

Contrary to our expectations, learning-related changes in PFC activation were comparable between adults and children. One interesting possibility is that motor sequence learning likely resembles action sequences and learning situations frequently encountered by children this age, thereby facilitating the implementation of familiar behavioral routines and the effective engagement of prefrontal control systems (Crone et al., 2006; Thomas et al., 2004). In contrast, paradigms with higher abstraction or complexity, such as working memory manipulation, response inhibition or task switching, often do reveal age-dependent differences in PFC recruitment (Crone et al., 2006; Rubia et al., 2006, 2007; Schwarze et al., 2023). Future studies incorporating younger age groups and systematically varying task demands as a function of children’s experience can help delineate the specific factors contributing to age-related differences in PFC activation during motor sequence learning.

It is also plausible that developmental changes in white matter microstructure contribute to improvements in motor skill learning by supporting faster processing speed and greater working memory capacity (Genc et al., 2023), underscoring the need for multimodal neuroimaging to elucidate how functional and structural brain changes interact to shape the maturation of cognitive control supporting motor sequence learning.

We observed less pronounced learning-related increases in M1 activation in children than in adults. Increasing M1 activation is considered to reflect the formation and storage of integrated, skill-dependent representations, that enable more efficient sequence execution (Hamano et al., 2021; Kornysheva & Diedrichsen, 2014). As learning progresses, M1, together with pre-SMA, is thought to support the transition from encoding discrete movements to encoding chunks, and ultimately to a more unified representation of the sequence (Lohse et al., 2014; Nambu et al., 2015; Penhune & Steele, 2012; Sakai et al., 2003; Verwey et al., 2002). From this perspective, the age-related differences in M1 activation in our study may indicate that adults were more effective at developing a unified representation of the sequence in M1, potentially through chunking, compared to children (Jones, 2012; Ruitenberg et al., 2013), leading to a greater increase in M1 activation (Penhune & Steele, 2012) and more efficient performance (Doyon et al., 2018; Ramkumar et al., 2016). This aligns well with our observation that, in adults, better task performance depended significantly on the effective recruitment of M1. This interpretation is further corroborated by the finding that adults showed a greater increase in representational similarity than children, suggesting more specialized and stable sequence encoding in M1 with practice (Huang et al., 2013; Lindenberger & Lövdén, 2019; Wiestler & Diedrichsen, 2013).

Interestingly, several recent studies have questioned the extent to which M1 represents sequences beyond individual movements (Berlot et al., 2020, 2021; Yokoi et al., 2018). Studies examining how sequence-related representations in M1 might differ between children and adults might therefore be particularly informative with regard to the nature of M1 representations. Furthermore, neural changes in M1 may reflect improved skill, but also differences in execution speed or movement variability (Berlot et al., 2020; Lutz et al., 2004; Orban et al., 2010). Adults in our sample performed faster and improved more than children, so group differences in M1 activation and representational stability could partly reflect age differences in performance. Recent work by Berlot et al. (2020) suggests, however, that variation in speed is unlikely to fully account for learning-related brain changes. When comparing paced and full-speed performance in adults, they found that overall activation and sequence-specific representations were largely preserved at matched speeds (Berlot et al., 2020). Their results imply that the increases in M1 activation and pattern stability observed in our study are unlikely to be a pure reflection of improvements in speed. We acknowledge that it is inherently difficult to disentangle learning from performance, because improvements in execution and speed are an integral part of learning. Nonetheless, future studies should include explicit comparisons between paced and full-speed performance to more fully dissociate effects of learning from effects of execution speed.

Beyond initial learning, consolidation processes facilitating skill maintenance and stabilization may differ between children and adults (Adi-Japha et al., 2019; Beck et al., 2024; Van Roy et al., 2024; Wilhelm et al., 2008). Wilhelm et al. (2008) found that in children, motor memory performance improved more during wakefulness than sleep whereas adults benefitted more from sleep-dependent consolidation. This highlights the need to examine how the age-related differences in M1 activation and representational stability observed in our study may interact with consolidation processes and affect performance in subsequent learning sessions, particularly during motor skill learning over multiple days.

In conclusion, our results advance the understanding of age-related differences in motor sequence learning by showing that, although M1 is engaged during motor sequence learning in middle childhood, its capacity to form stable, sequence-related neural representations is still maturing. This developmental limitation likely contributes to the more efficient motor sequence learning observed in adulthood.

## Materials and Methods

The hypotheses and methods of this study were pre-registered at the Open Science Framework (https://osf.io/gz36j).

### Participants

We recruited 57 children and 47 adults from the participant database at the Max Planck Institute for Human Development in Berlin, Germany. Eight adults and 18 children were excluded due to technical issues (three adults, one child), incomplete data (three adults, 16 children), left-handedness (one adult), or claustrophobia (one adult, one child). Our final sample for behavioral analyses included 39 children (17 female), aged between 7 and 10 years (*M* = 9.55; *SD* = 1.05), and 39 adults (19 female), aged between 20 and 32 years (*M* = 25.8; *SD* = 3.55). We deviated from our preregistration, which specified that we would recruit 7–8-year-olds, by including children up to 10 years to facilitate participant recruitment and ensure an adequate sample size. All participants were right-handed (Oldfield, 1971), native German speakers (for children, at least one parent was a native German speaker), had normal or corrected-to-normal vision, and no history of psychiatric or neurological disorders. The ethics committee of Freie Universität Berlin approved the study, which conformed to the guidelines of the Declaration of Helsinki (2008). We collected written informed consent from adult participants and children’s parents or legal guardians. Additionally, children gave verbal and written consent to participate.

### Experimental Design

The experimental design is depicted in Figure 9A. We tested participants on two consecutive days. On Day 1, participants completed several covariate measures, including a handedness test, working memory, and reaction time tasks. They then received instructions and practiced the main motor sequence learning task on a computer. Following practice, all participants performed one block of the motor sequence learning task, comprising two motor sequences, in an MRI simulator (i.e., identical to the MRI scanner, but without the magnetic field). Approximately 24 hours later, on Day 2, participants performed the motor sequence learning task in the MRI scanner. In the first scanning block, participants performed the two sequences from Day 1. Blocks two and three on Day 2 each included two new sequences that participants had not encountered before.

**Figure 9.**
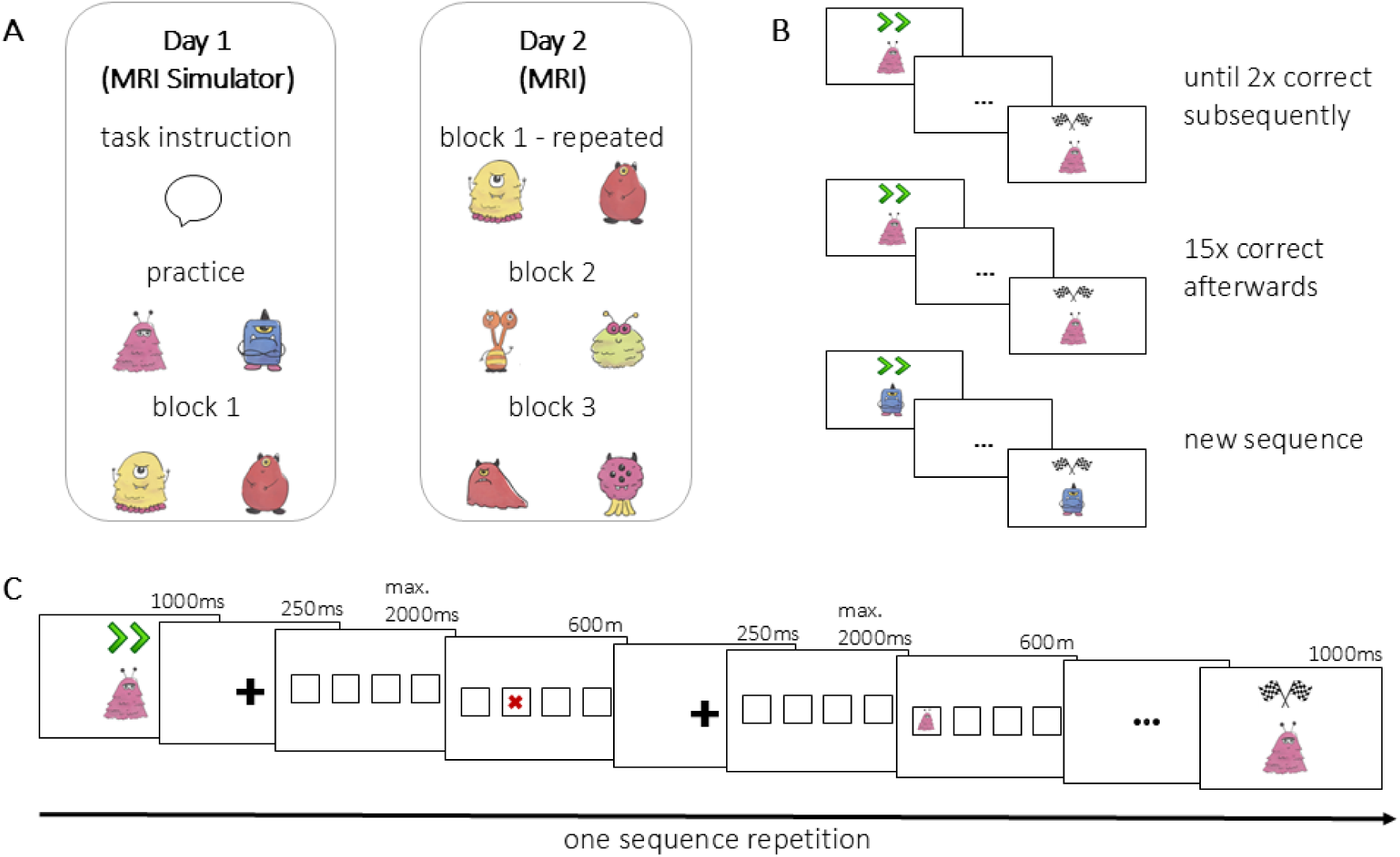
Experimental Design. ***A,*** Participants were tested on two consecutive days. On Day 1, participants were first instructed and then practiced the motor sequence learning task on a computer. Afterwards, they performed two sequences in an MRI simulator. On Day 2, six sequences were performed in the MRI scanner. The first two sequences were identical to the two sequences performed the day before in the MRI simulator. Thereafter, four new sequences were presented. ***B,*** To successfully perform the motor sequence learning task, participants had to execute two correct sequence repetitions consecutively. Afterwards, 15 correct sequence repetitions in total were necessary. This resulted in at least 17 correctly performed sequence repetitions for each participant, before the learning of a new sequence started. ***C,*** Example of one sequence repetition. Participants first saw the character with two green arrows displayed above it, indicating the start of the sequence. Subsequently, four white squares were displayed in a row corresponding to the four possible response buttons. After pressing one of the four buttons, participants received immediate feedback: a red cross appeared if they pressed the wrong button. If they responded correctly, the current character appeared in the selected square. When all elements of the sequence were pressed in the correct order, the character appeared with checkered flags over its head, indicating successful completion of one sequence repetition. *Characters in the figure were designed by Freepik* https://www.freepik.com*. In the original experiment, we used characters from Mr. Men Little Miss, which are not depicted here for copyright reasons*.

### Motor Sequence Learning Task

The motor sequence learning task in the present study was adapted from a study by Jenkins et al. (1994). We included a child-friendly cover story in which different characters were playing hide and seek (from https://www.mrmen.com/characters/). Participants were told to figure out the sequence of hideouts specific to each individual character. An example of one sequence repetition is presented in Figure 9C. In this associative visuomotor task, participants first saw one character with two green arrows displayed above it, indicating the start of the sequence. Subsequently, four white squares were displayed in a row at the center of the screen (for max. 2000 ms). Each square corresponded to one of four possible response buttons. Participants used their right index finger, middle finger, ring finger, and pinky finger, each corresponding to a single button (from left to right). Participants had to determine the button sequence associated with the initially shown character through trial and error. After pressing one of the four buttons, immediate feedback was provided for 600 ms: If the button press was incorrect, a red cross appeared in the selected square, and the participant continued to search for the correct button. If the button press was correct, the character appeared in the selected square. Next, the four white squares reappeared (max. 2000 ms), allowing the participant to select the next square via a button press. When all elements of the sequence were pressed in the correct order, the character appeared with checkered flags over its head (for 1000 ms), indicating successful completion of one sequence repetition. Between sequence repetitions, there was a jittered baseline period of 2000–8000 ms during which a fixation cross was presented.

To complete the learning of one sequence, participants had to fulfill two requirements (Figure 9B). First, they had to execute two consecutive correct sequence repetitions. Second, 15 correct sequence repetitions had to be completed following the two consecutive correct repetitions of the sequence. These 15 sequence repetitions did not need to be completed consecutively, i.e., they could be interleaved with erroneous repetitions. While the task specified a fixed criterion of at least 17 correct repetitions per sequence, the total number of sequence repetitions (including erroneous repetitions and any additional correct repetitions before reaching the first criterion of two subsequent correct repetitions) varied across participants. The completion of one sequence (i.e., after the predefined 17 correct sequence repetitions) was followed by an extended baseline fixation cross period (30 s) before the learning of the next sequence started.

Each block consisted of a short sequence and a long sequence. The short sequences consisted of four elements for children and six elements for adults, and the long sequences consisted of six elements for children and eight elements for adults. We determined these sequence lengths in behavioral pilots to ensure they were of comparable difficulty for the respective age group. The sequences and the order in which they were presented were identical for all participants within one age group. The six-element sequences (long for children, short for adults) were identical across age groups. Due to individual differences in learning and varying sequence lengths, the duration of one sequence repetition varied (children: short (4 elements), *M* = 6.34 s, *SD* = 2.26 s; long (6 elements), *M* = 9.50 s, *SD* = 3.23 s; adults: short (6 elements), *M* = 8.10 s, *SD* = 3.05 s; long (8 elements), *M* = 11.00 s, *SD* = 3.66 s). If a single block consisting of two sequences exceeded 20 minutes, the block was aborted (*n* = 5 children) and incompletely learned sequences were not included in the analyses. On average, children and adults did not differ in the duration of the scanning time for the execution of all repetitions of a sequence (children: *M* = 333.84 s, *SD* = 76.4 s; adults: *M* = 322.47 s, *SD* = 40.63 s; *t(48.71)* = 0.78, *p* = 0.44).

### Behavioral Analysis

We performed all analyses in R 4.2.2 (R Core Team, 2022) using RStudio (RStudio Team, 2020) with the tidyverse (Wickham et al., 2019), lme4 (Bates et al., 2015), lmerTest (Kuznetsova et al., 2017), performance (Lüdecke et al., 2021), and emmeans (Lenth, 2022) packages.

### Error Rates

To verify that the experimental manipulation was successful, and participants did indeed learn the sequences, we first divided sequence repetitions into two learning phases based on the criterion of two correct sequence repetitions in a row. The early learning phase was defined as all sequence repetitions performed until the sequence was correctly executed twice consecutively, including those two subsequent correct sequence repetitions. The late learning phase was defined as all sequence repetitions after the two correct sequence repetitions (15 correct repetitions for each participant plus an individual number of erroneous repetitions). As a result of this definition, the number of sequence repetitions in the two learning phases varied for each participant. However, the experimental procedure ensured that all participants had at least 17 correct repetitions for each sequence in addition to varying numbers of incorrect executions of each sequence.

We measured error rates in each participant for every sequence repetition as the number of errors in relation to the total amount of button presses made within that sequence repetition. Error rates were averaged across all repetitions of a sequence in each of the early and the late learning phases, separately for each sequence. Differences in error rates between the two learning phases were tested within each age group with paired t-tests, collapsing across short and long sequences. Note that we deviated from the ANOVA specified in the preregistration here due to violation of the normality assumption. We collapsed across all six sequences performed on Day 2 and did not distinguish between previously encountered and new sequences. This approach was applied in all subsequently reported analyses.

#### Response times

We computed mean response times (RTs) across the correct button presses for each sequence repetition in each participant. We determined RT outliers within each age group across sequences and repetitions. Sequence repetitions with RT values exceeding 3.29 standard deviations (*p* < .001, two-tailed; Tabachnick & Fidell, 2007) were excluded from subsequent analyses (1% of data). Due to a violation of the normality assumption in both age groups, RT data were log-transformed using the natural logarithm. The log-transformed RT data were normally distributed based on histograms and quantile–quantile plots (Q-Q plots).

We used linear mixed-effects models to examine performance improvements with learning across age groups. First, we compared age groups on the 6-element sequences that were identical in both age groups. To do so, we predicted log-transformed RTs by the fixed effects of age group (children vs. adults), the linear and quadratic effects of sequence repetition, and the two two-way interactions, linear sequence repetition × age group and quadratic sequence repetition × age group. The model also included random intercepts for participant and sequence. Second, we took the different sequence lengths incorporating age-matched difficulty into account. To this end, we fitted a linear mixed-effects model with log-transformed RTs as the dependent variable and fixed effects of age group (children vs. adults), sequence length (short vs. long) and the linear and quadratic effects of sequence repetition. We also included the two three-way interactions, linear sequence repetition × age group × sequence length and quadratic sequence repetition × age group × sequence length. Random intercepts for participant and sequence were also included in the model. We compared this model with simpler models that included only main effects and two-way interactions, using likelihood ratio tests. The model comparisons revealed that the inclusion of three-way interactions was unwarranted (all *p*s > 0.05). We also compared models with different parameterizations of repetition (linear only, linear plus quadratic, and a logarithmic term), and found that the model including both linear and quadratic repetition terms provided the best fit, likely reflecting the sparsely sampled later repetitions (i.e., high repetition numbers were present only for a small number of participants). Below, we report the best fitting model that included all main effects along with interactions between linear sequence repetition × age group, quadratic sequence repetition × age group, age group × sequence length, and quadratic effect sequence repetition × sequence length.

To account for a potential speed-accuracy trade-off, we performed control analyses in which we calculated inverse efficiency scores (Bruyer & Brysbaert, 2011). The inverse efficiency score combines speed and error into a single value calculated as 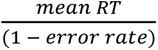 (Townsend & Ashby, 1983). This calculation penalizes RTs when committing errors. We analyzed the inverse efficiency score by fitting a linear mixed-effects model with the log-transformed inverse efficiency score as the dependent variable, using the same factors as described above for the RTs.

To limit the influence of single observations on the estimated effects, we removed all sequence repetitions beyond 42, corresponding to the maximum number of repetitions achieved in the adult group. A total of seven children had more than 42 repetitions of a single sequence resulting in the removal of 0.5% of data. Control analyses including all sequence repetitions for all participants did not change any of the results reported below.

### fMRI data acquisition and preprocessing

MR images were acquired in a supine position utilizing a CE certified 3-Tesla TIM Trio MRI scanner system (Siemens Medical Systems, Erlangen) with a 32-channel head coil. Whole-brain images were obtained for assessment of task-related activation with a T2*-weighted single-shot echo-planar imaging (EPI) sequence sensitive to BOLD contrast. The parameters utilized were: TR = 2000 ms, TE = 30 ms, flip angle = 80°, distance factor = 20%, isotropic voxel size = 3 mm^3^, 36 slices, FOV = 216 × 216 × 129 mm^3^, using a GRAPPA acceleration factor of 2. Slices were acquired in an interleaved fashion and aligned to the genu splenium of the corpus callosum. Structural images were acquired for co-registration of functional images using a three-dimensional T1-weighted magnetization prepared gradient-echo sequence (MPRAGE) with these parameters: repetition time (TR) = 2500 ms, echo time (TE) = 3.69 ms, inversion time (TI) = 1100 ms, flip angle = 7°, bandwidth = 200 Hz/pixel, acquisition matrix = 256 mm × 256 mm × 192 mm, isotropic voxel size = 1 mm^3,^ using a GRAPPA acceleration factor of 2; acquisition time = 5.58 min.

Data preprocessing and analysis of fMRI data were conducted using MRIQC version 0.15.1 (Esteban et al., 2017), fMRIPrep version 20.1.1 (Esteban et al., 2019, 2020), and SPM12 v7771 (https://www.fil.ion.ucl.ac.uk/spm/software/spm12) in MATLAB version 9.7.01190202 (MATLAB 2019b, The MathWorks, Inc., Natick, Massachusetts, USA). The MarsBaR toolbox version 0.44 (marsbar.sourceforge.net) was used for data extraction of relevant clusters. The xjView toolbox version 10.0 (https://www.alivelearn.net/xjview) and MRIcron v1.0.20190902 (https://www.nitrc.org/projects/mricron) were used to display results and report the anatomical locations of voxel clusters.

A total of five children were excluded from analyses because they had at least one run with more than 30% of TRs exceeding a framewise displacement of 0.4 mm (Dosenbach et al., 2017; Power et al., 2012), resulting in 34 child datasets (16 female) and 39 adult datasets available for fMRI analyses. Note that we deviated from the initial preregistration, where we specified exclusion based only on average absolute motion exceeding 3 mm, and used a more stringent threshold in the present analyses. We further deviated from the preregistration, stating preprocessing in SPM, and instead used fMRIPrep for preprocessing as it provides superior control for participant motion. We ran fMRIPrep using default preprocessing steps, including slice time correction, realignment, co-registration, and normalization to standard space using the MNI152NLin6ASym template. Subsequently, images were smoothed with an 8 mm full-width-half-maximum Gaussian kernel in SPM prior to analyses.

### General Linear Model and Region of Interest (ROI) Analysis

For each participant, a general linear model (GLM) was estimated to investigate whole-brain differences in task-related brain activation associated with the early and late phases of motor sequence learning as defined above. Each motor sequence was modeled with two regressors, one for sequence repetitions within the early and one for repetitions within the late learning phase, resulting in 12 regressors based on 6 unique sequences. Each sequence repetition was coded as an event with a duration equal to the time taken to complete the sequence repetition and convolved with a canonical hemodynamic response function (HRF). In addition, each sequence included a regressor of no interest, which incorporated feedback presentation, to exclude potential reward-related brain activations. To account for head motion artifacts, we included 31 regressors of no interest: 24 head motion parameters (6 base motion parameters, 6 temporal derivatives of 6 motion parameters, 12 quadratic terms of 6 motion parameters and their 6 temporal derivatives), framewise displacement per frame (Power et al., 2012), and the first six anatomical CompCor components estimated during preprocessing. CompCor uses a principal-component analysis approach to identify patterns of physiological noise and the inclusion of the components aids in the removal of noise from fMRI data (Behzadi et al., 2007). In addition, low-frequency drifts were removed using a 128 s high-pass filter. The first four volumes of each run were discarded to ensure stabilization of the magnetic resonance signal.

To identify the brain regions supporting motor skill learning in each of the learning phases, we computed the contrasts early > late learning phase and late > early learning phase at the whole-brain level, collapsing across age groups and sequence lengths.

Additionally, whole-brain two-sample t-tests comparing children and adults in each contrast were performed to examine brain regions showing potential age group differences in activation between the early and the late learning phases. All whole-brain analyses were corrected at *ppeak* < 0.05 FWE-corrected, *k* = 50 voxels.

Note that our preregistration primarily specified the whole-brain analyses described above. We extended this by implementing a more fine-grained ROI-based analysis, which allowed a more detailed examination of changes over time across individual sequence repetitions, rather than only comparing two discrete learning phases.

To define regions of interest (ROIs) based on our hypotheses, we took the significant clusters from the above whole-brain contrasts (collapsed across age groups and sequence lengths) and masked them with left and right inferior frontal gyrus (IFG), pars triangularis, and pars opercularis from the Harvard-Oxford atlas thresholded at 25% (HOCPAL; Desikan et al., 2006), and the left precentral gyrus and bilateral supplementary motor area as well as left and right putamen of the automated anatomical labelling atlas 3 (AAL3v1; Rolls et al., 2020).

We then estimated a GLM for each participant to examine brain activation during single sequence repetitions in the course of motor sequence learning. Each motor sequence was modeled with 17 regressors corresponding to the first 17 repetitions of that specific sequence, irrespective of whether errors occurred within the repetition. Additionally, we added a regressor of no interest, which incorporated remaining sequence repetitions, and the same regressors of no interest as the ones for the GLM above. For each ROI, parameter estimates were extracted from single sequence repetitions for each participant (Brett et al., 2002). Within each region and age group, cases with parameter estimate values exceeding 3.29 standard deviations (*p* < .001, two-tailed) were excluded from the ROI analyses (0.22% of data). For each region, a linear mixed-effects model was fitted with parameter estimates as the dependent variable and the fixed effects of age group (children vs. adults), linear sequence repetition, and quadratic sequence repetition (i.e., consistent with the slowing down in learning observed behaviorally). We also included interactions of both linear and quadratic sequence repetition with age group. All models included a random intercept per participant. In model testing, we first removed the quadratic sequence repetition × age group interaction and finally tested a model including only main effects. The best fitting model in each region was determined by likelihood ratio tests and is reported in the results.

In exploratory analyses relating differences in brain activation to differences in task performance, we performed a median split of RTs within each age group. Specifically, RTs were averaged over all sequence repetitions and sequences for each participant, and the median of these mean RTs was computed within each age group. This resulted in two performance subgroups: “high performers” (overall reaction times below the age group median) and “low performers” (overall reaction times above the age group median). This division into performance subgroups was subsequently incorporated as an additional factor in the linear mixed-effects model for each ROI.

### Representational Similarity Analysis (RSA)

All analysis scripts were written in Python v3.8.6 (Python Software Foundation, https://www.python.org/). A first-level analysis was conducted on the preprocessed task data using the Nipype (version 1.6.0; Gorgolewski et al., 2011) interface to FSL FEAT (using FSL 5.0.9; Jenkinson et al., 2012). The following events were modeled in the GLM for each sequence within each participant: 17 correct motor sequence repetitions (including the 2 consecutive correct repetitions that marked the end of the early learning phase, plus the 15 correct repetitions in the late learning phase); incorrect sequence repetitions; single correct sequence repetitions in the early phase before reaching 2 consecutive correct repetitions; and nuisance events corresponding to the indication of the start and end of a given sequence and the feedback presentation. In addition, we included motion parameters (rotation and translation parameters), framewise displacement per frame and the first six aCompCor components as nuisance regressors, identical to the GLMs described above. The first four volumes of each run were removed using Nibabel version 4.0.1 (Brett et al., 2022). No smoothing of the BOLD data was performed. We extracted the activation patterns for individual trials using a least squares separate (LSS) approach (Mumford et al., 2012).

Representational similarity analyses were performed using Nilearn version 0.9.1 (Abraham et al., 2014) within each participant separately. Based on previous research (Dayan & Cohen, 2011; Hamano et al., 2021; Penhune & Steele, 2012; Wiestler & Diedrichsen, 2013), the analyses focused on the left precentral gyrus ROI defined from the univariate analyses as described above. For each motor sequence, the extracted patterns of activation for each of the 17 correct sequence repetitions were first masked using the left precentral gyrus ROI and then merged to form a signal matrix. Thus, for each motor sequence we obtained a matrix with 17 rows corresponding to the minimum number of correct sequence repetitions and as many columns as the number of voxels *v* included in the ROI mask of interest. Using the signal matrix *SM*, we computed a representational similarity matrix (*RSM*) for each motor sequence *S*. The *RSM* is a correlation matrix where element (*i,j*) corresponds to the Pearson correlation coefficient between repetition *i* of motor sequence *S* and repetition *j* of motor sequence *S*.

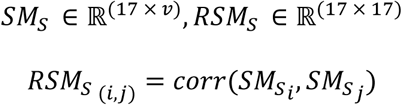

Correlation values of the resulting *RSM* were Fisher Z-transformed, and correlations between consecutive correct repetitions of the same motor sequence were extracted for each participant and each sequence, thereby operationalizing representational stability (i.e., pattern stability within a sequence).

A total of three runs (originating from one adult and two children) were excluded from the analyses due to technical issues. Correlation values exceeding 3.29 standard deviations (*p* < 0.001, two-tailed) within each age group were excluded from analyses (0.2% of data). We used mixed-effects models to examine changes in representational similarity with sequence repetition in left precentral gyrus. To this end, we tested for linear and quadratic effects of repetition pair, age group, and the interactions between these predictors, including random intercepts for participants and sequences. We compared this model to simpler models with only main effects and two-way interactions using likelihood ratio tests. The model comparisons revealed that the two-way interaction between the quadratic effect of repetition pair and age group was not warranted (all *p*s > .05). The best-fitting model reported below included the main effects of the linear and quadratic terms for repetition pair and age group. It also incorporated the interaction between the linear effect of repetition pair and age group.

### Psychophysiological Interaction Analysis (PPI)

We further conducted exploratory analyses examining how functional connectivity between PFC and M1 changed with sequence repetition. To this end, we used the left and right IFG and left precentral gyrus as ROIs, as defined above. To investigate how functional connectivity changed between ROIs with sequence repetition, we performed generalized psychophysiological interaction (gPPI) analyses using the CONN toolbox (v22; Whitfield-Gabrieli & Nieto-Castanon, 2012). The ROIs served once as a seed and once as a target region (i.e., left IFG–left M1, left M1–left IFG, right IFG–left M1, left M1–right IFG). In this ROI-to-ROI gPPI approach, the mean BOLD time series from each seed ROI and the task regressors corresponding to the first 17 sequence repetitions were entered into the model, along with their interaction terms. The interaction terms were computed by multiplying the seed ROI time series with each of the 17 task regressors. For each participant, a separate multiple regression model was estimated for each target ROI, including all 17 task regressors, the seed ROI time series, and the 17 interaction terms as predictors. The resulting gPPI estimates reflected changes in connectivity strength between each pair of the seed and target ROIs across repetitions of the motor sequence. Models were estimated using the unsmoothed preprocessed fMRI data, after running CONN’s default denoising pipeline including linear regression of potential confounding effects in the BOLD signal and temporal band-pass filtering. First-level analyses produced z-transformed correlation coefficients for each pair of ROIs per sequence repetition and participant, which were then used in subsequent linear mixed-effects models.

We excluded cases in each region pair and age group with connectivity estimate values exceeding 3.29 standard deviations (*p* < 0.001, two-tailed; 0.55% of data). To test whether the pairwise assignments yielded systematically different connectivity estimates, we used linear mixed-effects models with assignment (seed vs. target) as an additional fixed effect. As no significant differences were found, the connectivity estimates were averaged within each region pair for further analyses. We used the same procedure to test for differences between hemispheres (left vs. right). As there were no significant hemisphere differences, we averaged the connectivity estimates across hemispheres. The resulting hemisphere-averaged estimates were then used to examine how functional connectivity between PFC and M1 changed with sequence repetition. Using connectivity estimates as dependent variable, we tested for the fixed effects of linear and quadratic effects of sequence repetition, age group, and the interaction between these factors, including random intercepts for participants. The models were compared to simpler models using likelihood ratio tests, and the best fitting model included the main effects of linear and quadratic sequence repetition and age group.

## Acknowledgments

We acknowledge the financial support of the Max Planck Institute for Human Development. This work was conducted at the Max Planck Dahlem Campus of Cognition (MPDCC) of the Max Planck Institute for Human Development, Berlin, Germany. During the work on her dissertation, M.H. was a pre-doctoral fellow of the International Max Planck Research School on the Life Course (IMPRS LIFE, www.imprs-life.mpg.de; participating institutions: Max Planck Institute for Human Development, Freie Universität Berlin, Humboldt-Universität zu Berlin, University of Michigan, University of Virginia, University of Zurich). Y.F.’s work was supported by a research fellowship from the Jacobs Foundation (2023-1510-00). The authors thank Sara Bonati and Theodoros Koustakas for support with the RSA analyses and Julia Delius for editorial assistance.

## References

Abraham, A., Pedregosa, F., Eickenberg, M., Gervais, P., Mueller, A., Kossaifi, J., Gramfort, A., Thirion, B., & Varoquaux, G. (2014). Machine learning for neuroimaging with scikit-learn. Frontiers in Neuroinformatics, 8, Article 14.

Adi-Japha, E., Berke, R., Shaya, N., & Julius, M. S. (2019). Different post-training processes in children’s and adults’ motor skill learning. PLOS ONE, 14(1), Article e0210658. 10.1371/journal.pone.0210658

Barnea-Goraly, N., Menon, V., Eckert, M., Tamm, L., Bammer, R., Karchemskiy, A., Dant, C. C., & Reiss, A. L. (2005). White matter development during childhood and adolescence: A cross-sectional diffusion tensor imaging study. Cerebral Cortex, 15(12), 1848–1854. 10.1093/cercor/bhi062

Bassett, D. S., Yang, M., Wymbs, N. F., & Grafton, S. T. (2015). Learning-induced autonomy of sensorimotor systems. Nature Neuroscience, 18(5), 744–751. 10.1038/nn.3993

Bates, D., Mächler, M., Bolker, B., & Walker, S. (2015). Fitting linear mixed-effects models using lme4. Journal of Statistical Software, 67(1), 1–48. 10.18637/jss.v067.i01

Beck, M. M., Kristensen, F. T., Abrahamsen, G., Spedden, M. E., Christensen, M. S., & Lundbye-Jensen, J. (2024). Distinct mechanisms for online and offline motor skill learning across human development. Developmental Science, 27(6), Article e13536. 10.1111/desc.13536

Behzadi, Y., Restom, K., Liau, J., & Liu, T. T. (2007). A component based noise correction method (CompCor) for BOLD and perfusion based fMRI. NeuroImage, 37(1), 90–101. 10.1016/j.neuroimage.2007.04.042

Berlot, E., Popp, N. J., & Diedrichsen, J. (2020). A critical re-evaluation of fMRI signatures of motor sequence learning. eLife, 9, Article e55241.

Berlot, E., Popp, N. J., Grafton, S. T., & Diedrichsen, J. (2021). Combining repetition suppression and pattern analysis provides new insights into the role of M1 and parietal areas in skilled sequential actions. Journal of Neuroscience, 41(36), 7649–7661. 10.1523/JNEUROSCI.0863-21.2021

Brett, M., Anton, J.-L., Valabregue, R., Poline, J.-B., & others. (2002). Region of interest analysis using an SPM toolbox. 8th International Conference on Functional Mapping of the Human Brain, 16(2), 497.

Brett, M., Markiewicz, C. J., Hanke, M., Côté, M.-A., Cipollini, B., McCarthy, P., Jarecka, D., Cheng, C. P., Halchenko, Y. O., Cottaar, M., Larson, E., Ghosh, S., Wassermann, D., Gerhard, S., Lee, G. R., Wang, H.-T., Kastman, E., Kaczmarzyk, J., Guidotti, R., … freec84. (2022). Nipy/nibabel: [Computer software]. Zenodo. 10.5281/zenodo.6658382

Bruyer, R., & Brysbaert, M. (2011). Combining speed and accuracy in cognitive psychology: Is the inverse efficiency score (IES) a better dependent variable than the mean reaction time (RT) and the percentage of errors (PE)? Psychologica Belgica, 51(1), 5–13. 10.5334/pb-51-1-5

Casey, B. J., Tottenham, N., Liston, C., & Durston, S. (2005). Imaging the developing brain: What have we learned about cognitive development? Trends in Cognitive Sciences, 9(3), 104–110. 10.1016/j.tics.2005.01.011

Chein, J. M., & Schneider, W. (2012). The brain’s learning and control architecture. Current Directions in Psychological Science, 21(2), 78–84. 10.1177/0963721411434977

Clegg, B. A., DiGirolamo, G. J., & Keele, S. W. (1998). Sequence learning. Trends in Cognitive Sciences, 2(8), 275–281. 10.1016/S1364-6613(98)01202-9

Crone, E. A., Wendelken, C., Donohue, S., Van Leijenhorst, L., & Bunge, S. A. (2006). Neurocognitive development of the ability to manipulate information in working memory. Proceedings of the National Academy of Sciences USA, 103(24), 9315–9320. 10.1073/pnas.0510088103

Dayan, E., & Cohen, L. G. (2011). Neuroplasticity subserving motor skill learning. Neuron, 72(3), 443–454. 10.1016/j.neuron.2011.10.008

Desikan, R. S., Ségonne, F., Fischl, B., Quinn, B. T., Dickerson, B. C., Blacker, D., Buckner, R. L., Dale, A. M., Maguire, R. P., Hyman, B. T., Albert, M. S., & Killiany, R. J. (2006). An automated labeling system for subdividing the human cerebral cortex on MRI scans into gyral based regions of interest. NeuroImage, 31(3), 968–980. 10.1016/j.neuroimage.2006.01.021

Desimone, R., Duncan, J., & others. (1995). Neural mechanisms of selective visual attention. Annual Review of Neuroscience, 18(1), 193–222. 10.1146/annurev.ne.18.030195.001205

Dhawale, A. K., Smith, M. A., & Ölveczky, B. P. (2017). The Role of Variability in Motor Learning. Annual Review of Neuroscience, 40, 479–498. 10.1146/annurev-neuro-072116-031548

Diamond, A. (2002). Normal development of prefrontal cortex from birth to young adulthood: Cognitive functions, anatomy, and biochemistry. In D. T. Stuss & R. T. Knight (Eds.), Principles of frontal lobe function (pp. 466–503). Oxford University Press. 10.1093/acprof:oso/9780195134971.003.0029

Dosenbach, N. U. F., Koller, J. M., Earl, E. A., Miranda-Dominguez, O., Klein, R. L., Van, A. N., Snyder, A. Z., Nagel, B. J., Nigg, J. T., Nguyen, A. L., Wesevich, V., Greene, D. J., & Fair, D. A. (2017). Real-time motion analytics during brain MRI improve data quality and reduce costs. NeuroImage, 161, 80–93. 10.1016/j.neuroimage.2017.08.025

Doyon, J., Gabitov, E., Vahdat, S., Lungu, O., & Boutin, A. (2018). Current issues related to motor sequence learning in humans. Current Opinion in Behavioral Sciences, 20, 89–97. 10.1016/j.cobeha.2017.11.012

Esteban, O., Birman, D., Schaer, M., Koyejo, O. O., Poldrack, R. A., & Gorgolewski, K. J. (2017). MRIQC: Advancing the automatic prediction of image quality in MRI from unseen sites. PLOS ONE, 12(9), Article e0184661. 10.1371/journal.pone.0184661

Esteban, O., Ciric, R., Finc, K., Blair, R. W., Markiewicz, C. J., Moodie, C. A., Kent, J. D., Goncalves, M., DuPre, E., Gomez, D. E. P., Ye, Z., Salo, T., Valabregue, R., Amlien, I. K., Liem, F., Jacoby, N., Stojić, H., Cieslak, M., Urchs, S., … Gorgolewski, K. J. (2020). Analysis of task-based functional MRI data preprocessed with fMRIPrep. Nature Protocols, 15(7), 2186–2202. 10.1038/s41596-020-0327-3

Esteban, O., Markiewicz, C. J., Blair, R. W., Moodie, C. A., Isik, A. I., Erramuzpe, A., Kent, J. D., Goncalves, M., DuPre, E., Snyder, M., Oya, H., Ghosh, S. S., Wright, J., Durnez, J., Poldrack, R. A., & Gorgolewski, K. J. (2019). fMRIPrep: A robust preprocessing pipeline for functional MRI. Nature Methods, 16(1), 111–116. 10.1038/s41592-018-0235-4

Fandakova, Y., & Wenger, E. (2024). Chapter One - Skill learning in the developing brain: Interactions of control and representation systems. In K. D. Federmeier (Ed.), Psychology of learning and motivation (Vol. 81, pp. 1–40). Academic Press. 10.1016/bs.plm.2024.07.002

Fischer, S., Wilhelm, I., & Born, J. (2007). Developmental differences in sleep’s role for implicit off-line learning: Comparing children with adults. Journal of Cognitive Neuroscience, 19(2), 214–227. 10.1162/jocn.2007.19.2.214

Foerde, K., & Poldrack, R. A. (2009). Procedural learning in humans. In L. R. Squire (Ed.), Encyclopedia of neuroscience (pp. 1083–1091). Academic Press. 10.1016/B978-008045046-9.00783-X

Garner, K. G., & Dux, P. E. (2015). Training conquers multitasking costs by dividing task representations in the frontoparietal-subcortical system. Proceedings of the National Academy of Sciences, 112(46), 14372–14377. 10.1073/pnas.1511423112

Genc, S., Raven, E. P., Drakesmith, M., Blakemore, S.-J., & Jones, D. K. (2023). Novel insights into axon diameter and myelin content in late childhood and adolescence. Cerebral Cortex, 33(10), 6435–6448. 10.1093/cercor/bhac515

Gogtay, N., Giedd, J. N., Lusk, L., Hayashi, K. M., Greenstein, D., Vaituzis, A. C., Nugent, T. F., Herman, D. H., Clasen, L. S., Toga, A. W., Rapoport, J. L., & Thompson, P. M. (2004). Dynamic mapping of human cortical development during childhood through early adulthood. Proceedings of the National Academy of Sciences of the United States of America, 101(21), 8174–8179. 10.1073/pnas.0402680101

Gorgolewski, K., Burns, C., Madison, C., Clark, D., Halchenko, Y., Waskom, M., & Ghosh, S. (2011). Nipype: A flexible, lightweight and extensible neuroimaging data processing framework in python. Frontiers in Neuroinformatics, 5, Article 13.

Halsband, U., & Lange, R. K. (2006). Motor learning in man: A review of functional and clinical studies. Journal of Physiology, 99(4–6), 414–424. 10.1016/j.jphysparis.2006.03.007

Hamano, Y. H., Sugawara, S. K., Fukunaga, M., & Sadato, N. (2021). The integrative role of the M1 in motor sequence learning. Neuroscience Letters, 760, Article 136081. 10.1016/j.neulet.2021.136081

Hong, J.-Y., Gallanter, E., Müller-Oehring, E. M., & Schulte, T. (2019). Phases of procedural learning and memory: Characterisation with perceptual-motor sequence tasks. Journal of Cognitive Psychology, 31(5–6), 543–558. 10.1080/20445911.2019.1642897

Huang, Y., Zhen, Z., Song, Y., Zhu, Q., Wang, S., & Liu, J. (2013). Motor training increases the stability of activation patterns in the primary motor cortex. PLOS ONE, 8(1), Article e53555. 10.1371/journal.pone.0053555

Janacsek, K., Fiser, J., & Nemeth, D. (2012). The best time to acquire new skills: Age-related differences in implicit sequence learning across the human lifespan. Developmental Science, 15(4), 496–505. 10.1111/j.1467-7687.2012.01150.x

Jenkins, I. H., Brooks, D. J., Nixon, P. D., Frackowiak, R. S., & Passingham, R. E. (1994). Motor sequence learning: A study with positron emission tomography. Journal of Neuroscience, 14(6), 3775–3790. 10.1523/JNEUROSCI.14-06-03775.1994

Jenkinson, M., Beckmann, C. F., Behrens, T. E. J., Woolrich, M. W., & Smith, S. M. (2012). FSL. NeuroImage, 62(2), 782–790. 10.1016/j.neuroimage.2011.09.015

Jones, G. (2012). Why chunking should be considered as an explanation for developmental change before short-term memory capacity and processing speed. Frontiers in Psychology, 3, Article 167. 10.3389/fpsyg.2012.00167

Jongbloed-Pereboom, M., Nijhuis-van der Sanden, M. W. G., & Steenbergen, B. (2019). Explicit and implicit motor sequence learning in children and adults: The role of age and visual working memory. Human Movement Science, 64, 1–11. 10.1016/j.humov.2018.12.007

Karatekin, C., Marcus, D. J., & White, T. (2007). Oculomotor and manual indexes of incidental and intentional spatial sequence learning during middle childhood and adolescence. Journal of Experimental Child Psychology, 96(2), 107–130. 10.1016/j.jecp.2006.05.005

Karni, A., Meyer, G., Jezzard, P., Adams, M. M., Turner, R., & Ungerleider, L. G. (1995). Functional MRI evidence for adult motor cortex plasticity during motor skill learning. Nature, 377(6545), 155–158. 10.1038/377155a0

Karuza, E. A., Emberson, L. L., & Aslin, R. N. (2014). Combining fMRI and behavioral measures to examine the process of human learning. Neurobiology of Learning and Memory, 109, 193–206. 10.1016/j.nlm.2013.09.012

Koechlin, E., Ody, C., & Kouneiher, F. (2003). The architecture of cognitive control in the human prefrontal cortex. Science, 302(5648), 1181–1185. 10.1126/science.1088545

Kornysheva, K., & Diedrichsen, J. (2014). Human premotor areas parse sequences into their spatial and temporal features. eLife, 3, Article e03043. 10.7554/eLife.03043

Kuznetsova, A., Brockhoff, P. B., & Christensen, R. H. B. (2017). lmerTest package: Tests in linear mixed effects models. Journal of Statistical Software, 82(13). 10.18637/jss.v082.i13

Lenth, R. V. (2022). emmeans: Estimated marginal means, aka least-squares means. https://CRAN.R-project.org/package=emmeans

Lindenberger, U., & Lövdén, M. (2019). Brain plasticity in human lifespan development: The exploration–selection–refinement model. Annual Review of Developmental Psychology, 1(1), 197–222. 10.1146/annurev-devpsych-121318-085229

Lohse, K. R., Wadden, K., Boyd, L. A., & Hodges, N. J. (2014). Motor skill acquisition across short and long time scales: A meta-analysis of neuroimaging data. Neuropsychologia, 59(1), 130–141. 10.1016/j.neuropsychologia.2014.05.001

Lüdecke, D., Ben-Shachar, M. S., Patil, I., Waggoner, P., & Makowski, D. (2021). performance: An R package for assessment, comparison and testing of statistical models. Journal of Open Source Software, 6(60), Article 3139. 10.21105/joss.03139

Lukács, Á., & Kemény, F. (2015). Development of different forms of skill learning throughout the lifespan. Cognitive Science, 39(2), 383–404. 10.1111/cogs.12143

Lutz, K., Koeneke, S., Wüstenberg, T., & Jäncke, L. (2004). Asymmetry of cortical activation during maximum and convenient tapping speed. Neuroscience Letters, 373(1), 61–66. 10.1016/j.neulet.2004.09.058

Mayor-Dubois, C., Zesiger, P., Van der Linden, M., & Roulet-Perez, E. (2016). Procedural learning: A developmental study of motor sequence learning and probabilistic classification learning in school-aged children. Child Neuropsychology, 22(6), 718–734. 10.1080/09297049.2015.1058347

Meulemans, T., Van der Linden, M., & Perruchet, P. (1998). Implicit sequence learning in children. Journal of Experimental Child Psychology, 69(3), 199–221. 10.1006/jecp.1998.2442

Moore, D. R., Cowan, J. A., Riley, A., Edmondson-Jones, A. M., & Ferguson, M. A. (2011). Development of auditory processing in 6- to 11-yr-old children. Ear & Hearing, 32(3), 269–285. 10.1097/AUD.0b013e318201c468

Mumford, J. A., Turner, B. O., Ashby, F. G., & Poldrack, R. A. (2012). Deconvolving BOLD activation in event-related designs for multivoxel pattern classification analyses. NeuroImage, 59(3), 2636–2643. 10.1016/j.neuroimage.2011.08.076

Nambu, I., Hagura, N., Hirose, S., Wada, Y., Kawato, M., & Naito, E. (2015). Decoding sequential finger movements from preparatory activity in higher-order motor regions: A functional magnetic resonance imaging multi-voxel pattern analysis. European Journal of Neuroscience, 42(10), 2851–2859. 10.1111/ejn.13063

Nemeth, D., Janacsek, K., & Fiser, J. (2013). Age-dependent and coordinated shift in performance between implicit and explicit skill learning. Frontiers in Computational Neuroscience, 7, Article 147. 10.3389/fncom.2013.00147

Nguyen, Q. N., Michon, K. J., Vesia, M., & Lee, T. G. (2025). Dissociable causal roles of dorsolateral prefrontal cortex and primary motor cortex over the course of motor skill development. The Journal of Neuroscience, 45(20), Article e2015232025. 10.1523/JNEUROSCI.2015-23.2025

Oldfield, R. C. (1971). The assessment and analysis of handedness: The Edinburgh inventory. Neuropsychologia, 9(1), 97–113. 10.1016/0028-3932(71)90067-4

Olivo, G., Lövdén, M., Manzouri, A., Terlau, L., Jenner, B., Jafari, A., Petersson, S., Li, T.-Q., Fischer, H., & Månsson, K. N. T. (2022). Estimated gray matter volume rapidly changes after a short motor task. Cerebral Cortex, 32(19), 4356–4369. 10.1093/cercor/bhab488

Orban, P., Peigneux, P., Lungu, O., Albouy, G., Breton, E., Laberenne, F., Benali, H., Maquet, P., & Doyon, J. (2010). The multifaceted nature of the relationship between performance and brain activity in motor sequence learning. NeuroImage, 49(1), 694–702. 10.1016/j.neuroimage.2009.08.055

Penhune, V. B., & Steele, C. J. (2012). Parallel contributions of cerebellar, striatal and M1 mechanisms to motor sequence learning. Behavioural Brain Research, 226(2), 579–591. 10.1016/j.bbr.2011.09.044

Plebanek, D. J., & Sloutsky, V. M. (2017). Costs of selective attention: When children notice what adults miss. Psychological Science, 28(6), 723–732. 10.1177/0956797617693005

Poldrack, R. A. (2000). Imaging brain plasticity: Conceptual and methodological issues— A theoretical review. NeuroImage, 12(1), 1–13. 10.1006/nimg.2000.0596

Power, J. D., Barnes, K. A., Snyder, A. Z., Schlaggar, B. L., & Petersen, S. E. (2012). Spurious but systematic correlations in functional connectivity MRI networks arise from subject motion. NeuroImage, 59(3), 2142–2154. 10.1016/j.neuroimage.2011.10.018

R Core Team. (2022). *R: A language and environment for statistical computing*. R Foundation for Statistical Computing. https://www.R-project.org/

Ramkumar, P., Acuna, D. E., Berniker, M., Grafton, S. T., Turner, R. S., & Kording, K. P. (2016). Chunking as the result of an efficiency computation trade-off. Nature Communications, 7(1), Article 12176. 10.1038/ncomms12176

Rolls, E. T., Huang, C.-C., Lin, C.-P., Feng, J., & Joliot, M. (2020). Automated anatomical labelling atlas 3. NeuroImage, 206, Article 116189. 10.1016/j.neuroimage.2019.116189

RStudio Team. (2020). RStudio: Integrated development environment for R. RStudio, PBC. http://www.rstudio.com/

Rubia, K., Smith, A. B., Taylor, E., & Brammer, M. (2007). Linear age-correlated functional development of right inferior fronto-striato-cerebellar networks during response inhibition and anterior cingulate during error-related processes. Human Brain Mapping, 28(11), 1163–1177. 10.1002/hbm.20347

Rubia, K., Smith, A. B., Woolley, J., Nosarti, C., Heyman, I., Taylor, E., & Brammer, M. (2006). Progressive increase of frontostriatal brain activation from childhood to adulthood during event-related tasks of cognitive control. Human Brain Mapping, 27(12), 973–993. 10.1002/hbm.20237

Ruitenberg, M. F. L., Abrahamse, E. L., & Verwey, W. B. (2013). Sequential motor skill in preadolescent children: The development of automaticity. Journal of Experimental Child Psychology, 115(4), 607–623. 10.1016/j.jecp.2013.04.005

Sakai, K., Hikosaka, O., Miyauchi, S., Takino, R., Sasaki, Y., & Pütz, B. (1998). Transition of brain activation from frontal to parietal areas in visuomotor sequence learning. The Journal of Neuroscience, 18(5), 1827–1840. 10.1523/jneurosci.18-05-01827.1998

Sakai, K., Kitaguchi, K., & Hikosaka, O. (2003). Chunking during human visuomotor sequence learning. Experimental Brain Research, 152(2), 229–242. 10.1007/s00221-003-1548-8

Sanes, J. N., & Donoghue, J. P. (2000). Plasticity and primary motor cortex. Annual Review of Neuroscience, 23(1), 393–415. 10.1146/annurev.neuro.23.1.393

Savion-Lemieux, T., Bailey, J. A., & Penhune, V. B. (2009). Developmental contributions to motor sequence learning. Experimental Brain Research, 195(2), 293–306. 10.1007/s00221-009-1786-5

Schwarze, S. A., Laube, C., Khosravani, N., Lindenberger, U., Bunge, S. A., & Fandakova, Y. (2023). Does prefrontal connectivity during task switching help or hinder children’s performance? Developmental Cognitive Neuroscience, 60, Article 101217. 10.1016/j.dcn.2023.101217

Sydnor, V. J., Larsen, B., Bassett, D. S., Alexander-Bloch, A., Fair, D. A., Liston, C., Mackey, A. P., Milham, M. P., Pines, A., Roalf, D. R., Seidlitz, J., Xu, T., Raznahan, A., & Satterthwaite, T. D. (2021). Neurodevelopment of the association cortices: Patterns, mechanisms, and implications for psychopathology. Neuron, 109(18), 2820–2846. 10.1016/j.neuron.2021.06.016

Tabachnick, B. G., & Fidell, L. S. (2007). Using multivariate statistics (5th ed.). Allyn & Bacon/Pearson Education.

Tandoc, M. C., Nadendla, B., Pham, T., & Finn, A. S. (2024). Directing attention shapes learning in adults but not children. Psychological Science, 35(10), 1139–1154. 10.1177/09567976241263347

Taubert, M., Mehnert, J., Pleger, B., & Villringer, A. (2016). Rapid and specific gray matter changes in M1 induced by balance training. NeuroImage, 133, 399–407. 10.1016/j.neuroimage.2016.03.017

Thomas, K. M., Hunt, R. H., Vizueta, N., Sommer, T., Durston, S., Yang, Y., & Worden, M. S. (2004). Evidence of developmental differences in implicit sequence learning: An fMRI study of children and adults. Journal of Cognitive Neuroscience, 16(8), 1339–1351. 10.1162/0898929042304688

Toni, I., Krams, M., Turner, R., & Passingham, R. E. (1998). The time course of changes during motor sequence learning: A whole-brain fMRI study. NeuroImage, 8(1), 50–61. 10.1006/nimg.1998.0349

Townsend, J. T., & Ashby, F. G. (1983). Stochastic modeling of elementary psychological processes. CUP Archive.

Van Roy, A., Albouy, G., Burns, R. D., & King, B. R. (2024). Children exhibit a developmental advantage in the offline processing of a learned motor sequence. Communications Psychology, 2(1), Article 30. 10.1038/s44271-024-00082-9

Verwey, W. B., Lammens, R., & Honk, J. van. (2002). On the role of the SMA in the discrete sequence production task: A TMS study. Neuropsychologia, 40(8), 1268–1276. 10.1016/S0028-3932(01)00221-4

Weiermann, B., & Meier, B. (2012). Incidental sequence learning across the lifespan. Cognition, 123(3), 380–391. 10.1016/j.cognition.2012.02.010

Whitfield-Gabrieli, S., & Nieto-Castanon, A. (2012). *Conn*: A functional connectivity toolbox for correlated and anticorrelated brain networks. Brain Connectivity, 2(3), 125–141. 10.1089/brain.2012.0073

Wickham, H., Averick, M., Bryan, J., Chang, W., McGowan, L. D., François, R., Grolemund, G., Hayes, A., Henry, L., Hester, J., Kuhn, M., Pedersen, T. L., Miller, E., Bache, S. M., Müller, K., Ooms, J., Robinson, D., Seidel, D. P., Spinu, V., … Yutani, H. (2019). Welcome to the Tidyverse. Journal of Open Source Software, 4(43), Article 1686. 10.21105/joss.01686

Wiestler, T., & Diedrichsen, J. (2013). Skill learning strengthens cortical representations of motor sequences. eLife, 2, Article e00801. 10.7554/eLife.00801

Wilhelm, I., Diekelmann, S., & Born, J. (2008). Sleep in children improves memory performance on declarative but not procedural tasks. Learning & Memory, 15(5), 373–377. 10.1101/lm.803708

Wu, T., Kansaku, K., & Hallett, M. (2004). How self-initiated memorized movements become automatic: A functional MRI study. Journal of Neurophysiology, 91, 1690–1698. 10.1152/jn.01052.2003

Yokoi, A., Arbuckle, S. A., & Diedrichsen, J. (2018). The role of human primary motor cortex in the production of skilled finger sequences. Journal of Neuroscience, 38(6), 1430–1442. 10.1523/JNEUROSCI.2798-17.2017

Zwart, F. S., Vissers, C. Th. W. M., Kessels, R. P. C., & Maes, J. H. R. (2019). Procedural learning across the lifespan: A systematic review with implications for atypical development. Journal of Neuropsychology, 13(2), 149–182. 10.1111/jnp.12139

