## Supplementary Material for "Motor Sequence Learning in Children and Adults: Age Differences in the Time Course of Brain Activation and Representational Stability"

**Table S1**

*Linear Mixed-Effects Model of Raw Response Times for Six-Element Sequences*

| | Estimate | SE | p-value |
|---|---|---|---|
| Intercept | 311.80 | 15.72 | < 0.001 |
| repetition | -14.23 | 1.40 | < 0.001 |
| I(sequence repetition^2) | 0.48 | 0.06 | < 0.001 |
| age group | 110.41 | 21.80 | < 0.001 |
| sequence repetition × age group | 9.05 | 1.57 | < 0.001 |
| I(sequence repetition^2) × age group | -0.38 | 0.06 | < 0.001 |

*Note.* Results of the linear mixed-effects model with raw response times as the dependent variable and age group (children vs. adults) the linear and quadratic effects of sequence repetition and their interactions (linear sequence repetition × age group and quadratic sequence repetition × age group) as predictors.

**Table S2**

*Linear Mixed-Effects Model of Inverse Efficiency Scores for Six-Element Sequences*

| | Estimate | SE | p-value |
|---|---|---|---|
| Intercept | 6.08 | 0.05 | < 0.001 |
| repetition | -0.12 | 0.01 | < 0.001 |
| I(sequence repetition^2) | 0.00 | 0.00 | < 0.001 |
| age group | 0.24 | 0.07 | < 0.001 |
| sequence repetition × age group | 0.08 | 0.01 | < 0.001 |
| I(sequence repetition^2) × age group | -0.00 | 0.00 | < 0.001 |

*Note*. Results of the linear mixed-effects model with log-transformed inverse efficiency scores as the dependent variable and age group (children vs. adults), the linear and quadratic effects of sequence repetition, and their interactions (linear sequence repetition × age group and quadratic sequence repetition × age group) as predictors.

**Table S3**

*Linear Mixed-Effects Model of Raw Response Times for Both Sequence Lengths*

| | Estimate | SE | p-value |
|---|---|---|---|
| Intercept | 302.17 | 16.43 | < 0.001 |
| repetition | -11.98 | 0.72 | < 0.001 |
| I(sequence repetition^2) | 0.38 | 0.03 | < 0.001 |
| age group | 132.43 | 14.52 | < 0.001 |
| sequence length | 24.10 | 18.47 | 0.238 |
| sequence repetition × age group | 6.34 | 0.89 | < 0.001 |
| I(sequence repetition^2) × age group | -0.19 | 0.03 | < 0.001 |
| sequence length × age group | -33.88 | 4.60 | < 0.001 |
| I(sequence repetition^2) × sequence length | -0.08 | 0.01 | < 0.001 |

*Note.* Results of the linear mixed-effects model with raw response times as the dependent variable. Predictors were age group (children vs. adults), sequence length (short vs. long), the linear and quadratic effects of sequence repetition, as well as the interactions between the linear and quadratic effects of sequence repetition and age group, between age group and sequence length and between the quadratic effect of sequence repetition and sequence length.

**Table S4**

*Linear Mixed-Effects Model of Inverse Efficiency Scores for Both Sequence Lengths*

| | Estimate | SE | p-value |
|---|---|---|---|
| Intercept | 6.09 | 0.06 | < 0.001 |
| repetition | -0.10 | 0.00 | < 0.001 |
| I(sequence repetition^2) | 0.00 | 0.00 | < 0.001 |
| age group | 0.26 | 0.05 | < 0.001 |
| sequence length | 0.18 | 0.07 | 0.035 |
| sequence repetition × age group | 0.05 | 0.00 | < 0.001 |
| I(sequence repetition^2) × age group | -0.02 | 0.00 | < 0.001 |
| sequence length × age group | -0.05 | 0.18 | 0.005 |
| I(sequence repetition^2) × sequence length | -0.00 | 0.00 | < 0.001 |

*Note.* Results of the linear mixed-effects model with log-transformed inverse efficiency scores as the dependent variable. Predictors were age group (children vs. adults), sequence length (short vs. long), the linear and quadratic effects of sequence repetition, as well as the interactions between the linear and quadratic effects of sequence repetition and age group, between age group and sequence length and between the quadratic effect of sequence repetition and sequence length.

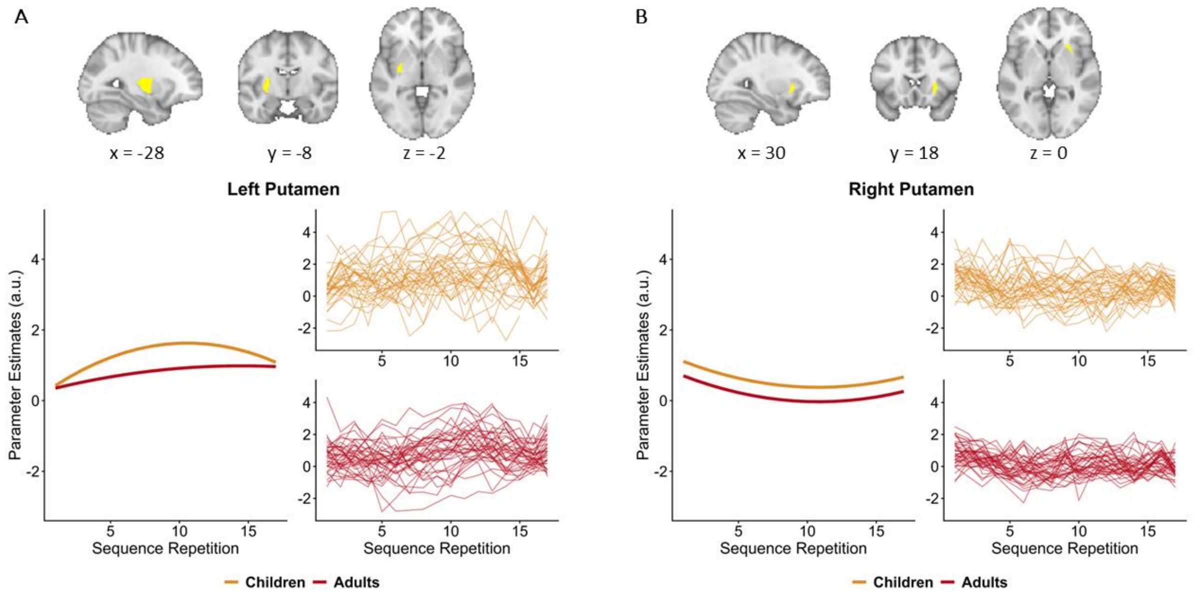


**Figure S1**
Brain activation in left and right putamen across sequence repetitions during learning. Model estimates for left (A) and right (B) putamen. For each ROI, the left panel depicts fixed-effect estimates from the linear mixed effect model testing for age-related differences in the linear and quadratic effects of sequence repetition. The right panel shows individual activation data for the first 17 sequence repetitions for children (in yellow) and adults (in red). In the left putamen, activation increased significantly across sequence repetitions. We observed a significant interaction between the linear and quadratic effects of sequence repetition and age group, suggesting that both the initial increase and the subsequent slowing in activation were more pronounced in children than in adults. In the right putamen, activation was overall higher in children than in adults. We observed a significant linear decrease with sequence repetition accompanied by a significant quadratic effect, indicating a deceleration of activation decreases over time. This pattern of changes was comparable across age groups.

**Table S5**
*Linear Mixed-Effects Model of Left Putamen Estimates*

| | Estimate | SE | p-value |
|---|---|---|---|
| Intercept | 0.16 | 0.18 | 0.38 |
| repetition | 0.28 | 0.03 | < 0.001 |
| I(sequence repetition^2) | -0.01 | 0.00 | < 0.001 |
| age group | 0.09 | 0.03 | 0.71 |
| sequence repetition × age group | 0.18 | 0.05 | < 0.001 |
| I(sequence repetition^2) × age group | 0.01 | 0.00 | < 0.001 |

*Note.* Results of a linear mixed-effects model with left putamen estimates as the dependent variable. Predictors were age group (children vs. adults), the linear and quadratic effects of sequence repetition, and their interactions (linear sequence repetition × age group and quadratic sequence repetition × age group).

**Table S6**

*Linear Mixed-Effects Model of Right Putamen Estimates*

| | Estimate | SE | p-value |
|---|---|---|---|
| Intercept | 1.27 | 0.1 | < 0.001 |
| repetition | -0.17 | 0.02 | < 0.001 |
| I(sequence repetition^2) | 0.01 | 0.00 | < 0.001 |
| age group | -0.41 | 0.09 | < 0.001 |

*Note.* Results of the linear mixed-effects model with right putamen parameter estimates as the dependent variable. Predictors were age group (children vs. adults) and the linear and quadratic effects of sequence repetition. Including interactions was not warranted.
